# Discovering the unseen: a performance comparison of taxonomic classification methods for unknown DNA barcodes

**DOI:** 10.1101/2025.10.13.681976

**Authors:** Johanna Orsholm, Alessandro Zito, Panu Somervuo, Jesse P Harrison, Markus Koskela, Otso Ovaskainen, Mariana P Braga, Nicolas Chazot, Tomas Roslin, Brendan Furneaux

## Abstract

1. DNA barcoding and metabarcoding have emerged as cost-efficient, standardized methods for characterizing local biodiversity. Based on the sequencing of a small targeted gene fragment, it is theoretically possible to identify a wide diversity of taxa by comparing them with reference sequence databases. However, a key challenge for accurate taxonomic classification is the incompleteness of such databases, leading to most query sequences lacking species-level matches.
2. Where species-level matches are missing, it may be possible to classify query sequences to a higher taxonomic level, such as genus or family, based on the similarity of related reference taxa. The challenge then lies in confidently recognizing whether the sequence belongs to an unobserved (here, “novel”) taxon on a given taxonomic level.
3. In this study, we evaluated the performance and utility of several methods for taxonomic classification. Methods were assessed based on the classification accuracy of both observed and novel taxa, training time, space requirements, and run time. We did this for two cases: the COI barcode for arthropods, and the ITS barcode for fungi, with the latter representing an instance with substantially greater sequence similarity variation within classes. To test classification of novel taxa, we used well-curated datasets with partially distinct taxonomic distribution between the training and test set. Novel taxa occurred at all evaluated taxonomic levels, such as novel species in observed genera and novel genera in observed families. We further assessed the effect on performance when shifting from full-length barcodes to shorter sequences as generated through metabarcoding in the testing dataset.
4. This study sheds light on the strengths and limitations of different classification algorithms across varied ecological contexts and provides guidance for researchers in selecting suitable algorithms for DNA barcoding and metabarcoding applications. In particular, it demonstrates the supreme performance of phylogenetic placement methods such as EPA-ng for classification of arthropod COI barcodes, and composition-based classifiers such as SINTAX, RDP-NBC, and IDTAXA for fungal ITS.

## 1 Introduction

During recent decades, DNA barcoding has emerged as a versatile method to characterize biological samples (Hebert et al., 2003). DNA barcoding relies on short, standardized gene segments to classify unknown specimens within a Linnaean taxonomy using previously assembled DNA reference databases. Such a procedure is especially effective when combined with high-throughput sequencing technologies, which enable both rapid parallel sequencing of a large number of individual specimens (Shokralla et al., 2014) and analysis of mixed samples in bulk, also known as DNA metabarcoding (Riaz et al., 2011). As a result, sequencing pipelines have become indispensable tools for large-scale biodiversity analysis and monitoring, enabling the identification of species across different taxa and allowing researchers to rapidly assess species diversity in various ecosystems (Gostel and Kress, 2022; Niskanen et al., 2023; Van Klink et al., 2022).

Early work confirming the utility of DNA barcodes for taxonomic classification of unknown specimens has typically focused on cases where there are close matches for the query sequences in the reference database; see Hebert et al. (2003); Schoch et al. (2012); Janzen et al. (2005); Deshpande et al. (2016). However, these ideal conditions are rare: many taxonomic branches from highly diverse groups are either undescribed or currently lack reference sequences, as evident from a striking difference between current estimates of global species richness and the actual sizes of the annotated libraries. For example, insect species diversity is estimated at 2.6-7.2 million (Stork, 2018), while well-curated reference databases account for roughly 1.2 million species-level sequence clusters with only ≈283 thousand annotated with formal species names (statistics downloaded on 2025-08-15 from https://www.boldsystems.org/). As for fungi, estimates oscillate between 1-12 million species, but databases present ≈170 thousand clusters with only ≈53 thousand formally annotated (Ratnasingham et al., 2024; Abarenkov et al., 2025, statistics downloaded on 2025-08-15 from https://unite.ut.ee/). Additionally, sequencing efforts have shown a clear geographic bias towards Europe and North America (Leandro et al., 2024; Khomich et al., 2018), leaving many of the most diverse regions of the world underrepresented.

Due to the limited coverage of diverse taxonomic groups and geographical regions, classifying specimens from taxa lacking reference sequences is a central challenge of DNA barcoding. We refer to such a category as the *novel taxa*, in contrast to *observed taxa* that are represented in the reference database. In this context, novel taxa thus include both known species lacking reference sequences and currently undescribed species. In a realistic scenario, a classification algorithm must accomplish two tasks: (1) it must predict if a query sequence belongs to an observed species and, if not, (2) it must correctly classify the higher taxonomic ranks of novel taxa, up to the last observed level; for instance, the family of novel genera, and the genus of novel species. The aim of this work is to test the ability of current taxonomic classifiers to satisfy both requirements.

When designing a taxonomic classification algorithm, one must first handle the biological heterogeneity of the commonly employed barcode regions. In principle, a genetic marker is suitable for taxonomic classification if the variation between species is larger than the variation within species (Somervuo et al., 2017; Hebert et al., 2004). These properties are seen in the mitochondrial cytochrome c oxidase subunit 1 (COI; Hebert et al., 2003) in animals, and in the nuclear ribosomal internal transcribed spacer (ITS; Schoch et al., 2012) in fungi. The traditional COI barcode region, known as the Folmer region, is usually 658 base pairs (bp) long (Folmer et al., 1994), and allows for relatively simple sequence alignments thanks to the strong functional constraints on its length and amino acid sequence (Pentinsaari et al., 2016). In contrast, the ITS barcode consists of two spacers—ITS1 and ITS2—surrounding the non-coding 5.8S rRNA, each varying widely in both base arrangement and sequence length (Schoch et al., 2012). As a consequence, it is practically impossible to produce reliable multiple sequence alignments across broad taxonomic groups (Lindahl et al., 2013). In fungi, ITS varies between 400 and 800 bp (Manter and Vivanco, 2007), but in some cases it may be up to 1600 bp in length (Feibelman et al., 1994). Hence, classifying aligned or non-aligned queries often requires distinct modeling assumptions and pre-processing steps, adding an extra layer of difficulty.

Over the past two decades, a wealth of algorithms have been developed to classify query sequences using taxonomically labeled reference sequences. Here, we test the performance of such taxonomic classification methods for DNA barcodes, specifically focusing on the issue of incomplete reference libraries. In contrast to previous studies, we use real-world geographical and methodological divisions to generate partially overlapping reference and test sequence sets. Combining historical methods with more novel approaches, we focus on stand-alone programs using a variety of algorithms either designed or commonly used for taxonomic classification of barcodes. We specifically consider two distinct biological cases: the COI barcode as a protein-coding gene with conserved length, and ITS as a non-coding sequence of variable length. In the case of COI, we also consider amino acid translations of the barcode for algorithms that accept protein sequences. Additionally, we evaluate performance at classifying a subregion of the query sequences. This comparison is motivated by practical considerations: due to the limitations on sequence length posed by some high-throughput sequencing methods, a shorter subregion of the markers is often targeted for DNA metabarcoding, resulting in around 400 bp for COI (Elbrecht et al., 2019) and 300-400 bp for ITS (Winand et al., 2025). Hence, our goal is to mimic the shorter metabarcoding reads, which are associated with lower sequencing costs. Our metrics of comparison are classification accuracy, calibration of confidence scores, prediction coverage, and use of computational resources. *A priori*, we expect (i) a trade-off between accuracy and coverage, and (ii) higher accuracy at the expense of higher computational costs for algorithms that rely on complex models of taxonomic structure in sequence data.

### 1.1 Overview of taxonomic classifiers

Taxonomic classification methods aim to label sequences using reference databases of DNA barcodes, often employing markedly different strategies such as similarity searches (Altschul et al., 1990; Camacho et al., 2009; Vu et al., 2022; Lanzen et al., 2012), probabilistic modeling (Zito et al., 2023; Somervuo et al., 2016) and phylogenetic inference (Barbera et al., 2019). However, no classification algorithm is universally superior to the others *a priori*. Rather, each method is originally designed for a specific purpose, either favoring speed of prediction, unbiased estimation of prediction probabilities, or the capacity to recognize novel taxa. Recently, Hleap et al. (2021) grouped taxonomic assignment algorithms into four categories: 1) similarity-based; 2) composition-based; 3) probabilistic; and 4) phylogenetic. We argue that recent advances in machine learning and deep learning methodologies have since led to the birth of a novel fifth category, that of 5) *neural network classifiers*. From each category, we selected representative methods and tested their performance; See Table 1 for a summary, and refer to Table S1 for additional algorithms and motivations for their exclusion from the comparison.

**Table 1:**
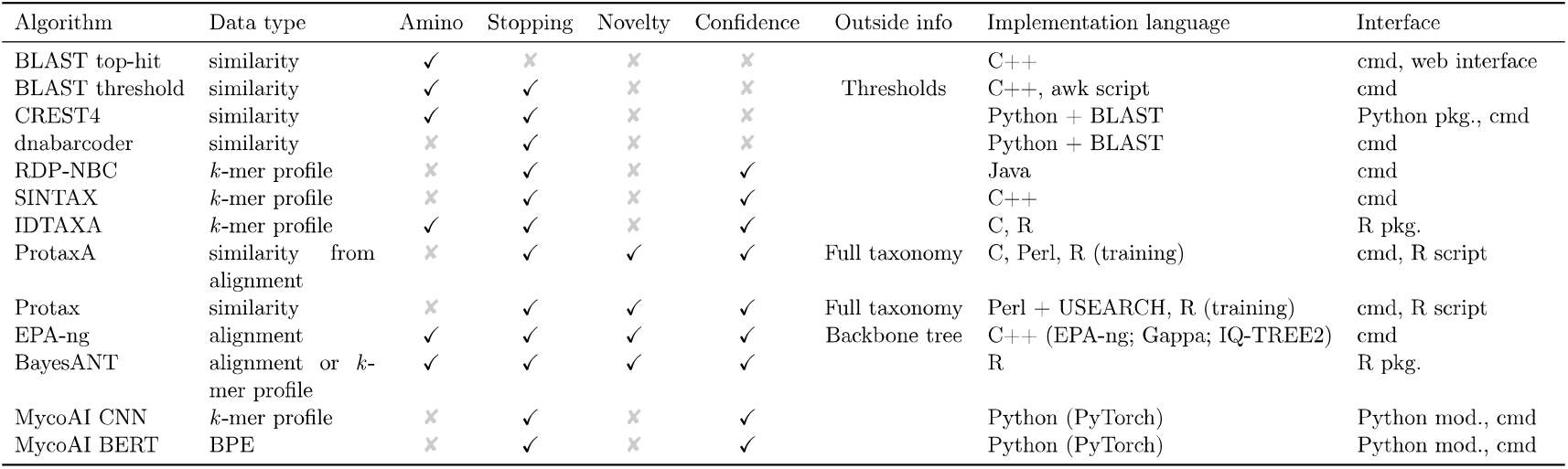
Features of selected taxonomic classification algorithms. Amino: All selected algorithms operate on nucleotide sequences; some can additionally be applied to amino acid sequences. Stopping: Does the algorithm have any mechanism to avoid over-classifying sequences? Novelty: Does the algorithm distinguish between low classification certainty for existing categories and prediction of new categories which were not present in the reference? Confidence: Does the algorithm produce a numerical score on a fixed scale (i.e., 0-1 or 0-100) for the confidence of its classifications at each rank? Outside info: Does the algorithm allow additional data beyond the sequences and taxonomic labels of the training data? If so, what? Implementation language: What language(s) is/are used for the implementation? Does it depend on any other software? Interface: How do you use it?

**Similarity-based classifiers** use pairwise alignment of query sequences to the reference sequences as the basis of classification. BLAST (Altschul et al., 1990; Camacho et al., 2009) is a commonly used tool for sequence similarity search. While it is not specifically designed for taxonomic classification, a widespread first approach is to simply take the taxonomy of the top BLAST hit; see Bonato et al. (2022); Vass et al. (2022) for examples. A slightly more sophisticated approach uses *a priori* similarity thresholds to establish whether to accept the top hit classification at each taxonomic rank (Tedersoo et al., 2021). Here, a similarity threshold of 97% has been widely used for species-level identification across different taxa and barcode markers (Tedersoo et al., 2022; Porter and Hajibabaei, 2020). These thresholds can be further optimized for different taxa, as in dnabarcoder (Vu et al., 2022), or replaced by instead considering multiple hits simultaneously, assigning the last common ancestor (LCA) of the *k* nearest neighbors (KNN) from a BLAST search, as in CREST4 (Lanzen et al., 2012). Thresholding and KNN-consensus both limit over-classification, which occurs when sequences from novel taxa are assigned taxonomic labels at ranks where no true match exists. The KNN-consensus method can also help avoid over-classification in cases where closely related species have identical barcode sequences. However, neither method provides a numerical confidence score or distinguishes between uncertain classifications and novel taxa.

**Composition-based classifiers** avoid the need for computationally expensive pairwise alignments by decomposing both queries and reference sequences into *k*-mers, i.e., distinct overlapping subsequences of length *k*. A sequence can then be described by its *k-*mer profile - a fixed-length vector with one entry for each possible *k-*mer. The length of the vector is 4^*k*^for nucleotide sequences and 20^*k*^ for amino acid sequences. From this category, we selected the RDP naïve Bayesian classifier (RDP-NBC, Wang et al., 2007), SINTAX (Edgar, 2016), and IDTAXA (Murali et al., 2018). All three models use bootstrap samples of the *k-*mer composition of the query sequences to produce numerical confidence scores for the classification at each taxonomic rank. This helps control over-classification in a more granular way than the similarity search methods, but does not directly address novelty in the taxonomy.

**Probabilistic classifiers** use a statistical framework to estimate the probability that a query sequence belongs to a given taxon at each taxonomic rank. From this category, we selected BayesANT (Zito et al., 2023) and PROTAX (Somervuo et al., 2016). PROTAX is a model framework that can incorporate any covariates as predictors of taxonomy (e.g., sequence similarity), allowing users to tailor the model to better fit their specific needs. There are also concrete implementations designed and trained for specific taxonomic groups, such as FinPROTAX for arthropods (Roslin et al., 2022) and Protax-Fungi for fungi (Abarenkov et al., 2018), which can be readily applied without further model customization. Instead, BayesANT leverages Bayesian nonparametric species sampling priors (Pitman, 1996; De Blasi et al., 2015) to model the taxonomic tree (Rigon et al., 2025). Each taxon and its frequency of appearance are assumed to be a realization from a rank-specific Pitman-Yor process (Pitman and Yor, 1997), which allows novel taxonomic nodes to appear. Hence, both BayesANT and PROTAX explicitly distinguish between classification uncertainty and novel taxa. Both have variants that can operate either on unaligned or globally aligned sequences, and while the latter may be both faster and higher-performing, it is limited to sequences with low length polymorphism, such as the COI barcode for animals. Moreover, BayesANT and PROTAX provide confidence scores designed to be interpreted as well-calibrated probabilities. In addition, PROTAX can leverage information from a reference taxonomy that includes taxa lacking associated reference sequences.

**Phylogenetic placement classifiers** place query sequences individually into an existing phylogenetic tree generated from a set of reference sequences. In this framework, novel taxa are cleanly represented as placement on branches outside the clades which represent observed taxa at the rank under consideration. Construction of the reference tree is left to the user, with a wide range of algorithms available but differing in speed and accuracy. There are several opportunities to incorporate additional information beyond the reference sequences at the stage of building the reference tree, including constraining the topology to conform to the taxonomic hierarchy or to published phylogenies. Alternatively, the reference tree can be generated using longer sequences or additional genes beyond the region covered by the query sequences. From this category we selected EPA-ng (Barbera et al., 2019), a maximum-likelihood based phylogenetic placement algorithm that calculates the likelihood of placement on each branch of the tree for each query sequence, according to a chosen evolutionary model. From these placements, Gappa (Czech et al., 2020), a helper tool for phylogenetic placement, can perform the mapping to taxonomic classification. Gappa can also report classification probabilities when they are provided by the phylogenetic placement algorithm. However, phylogenetic placement typically requires that the query and reference sequences are globally aligned, making it unsuitable for ITS fungal sequences.

**Neural network classifiers** are united by their use of a learned vector representation for barcode sequences. From this category, we selected MycoAI (Romeijn et al., 2024), which implements both a convolutional neural network (CNN) operating on sequences represented as real-valued *k-*mer profiles, and a bidirectional transformer (BERT) operating on byte-pair encoded sequences. Both architectures use hierarchical label smoothing during training and a multi-head output for the different taxonomic ranks in order to borrow information between taxonomic ranks and provide separate predictions and confidence values at each rank.

Previous comparisons of taxonomic classifiers have found that simple top-hit algorithms, such as BLAST, often achieve the highest classification accuracy of observed species (Edgar, 2018; Hleap et al., 2021), making them attractive choices for study systems with extensive reference libraries. However, classification accuracy typically declines with a decrease in the similarity between the query sequence and the top hit reference sequence (Edgar, 2018; Richardson et al., 2017), making it challenging to reliably classify novel taxa. As previously described, taxonomic classifiers should ideally make predictions down to the lowest observed rank of a query sequence. Thus, setting conservative confidence thresholds for predictions can help reduce misclassification and over-classification - but may simultaneously lead to high rates of under-classification, where too few taxonomic ranks are predicted. Classification of novel taxa is further complicated by the lack of a consistent relationship between sequence similarity and the lowest common rank across taxonomic branches (Edgar, 2018). The complexity of this task calls for well-calibrated estimations of prediction uncertainty (Somervuo et al., 2017), allowing researchers to make informed decisions about retaining classifications and potentially incorporating uncertainty in downstream analyses.

## 2 Materials and methods

### 2.1 Data description and pre-processing

In our evaluation, models were trained on a set of barcodes, called the *training set*, which had a partial taxonomic overlap with the sequences used to assess model performance, the *test set*. We also generated a *testshort set* in which a standardized subregion of the sequences in the test set was selected, to represent the shorter sequences that are often obtained with metabarcoding.

All sequences in both sets were associated with a species-level identification. However, in some cases higher ranks are missing from the classifications. These can occur for two reasons. First, the “secondary ranks” - in our COI dataset subfamily and tribe - are considered optional, and are only used in diverse groups where additional levels of classification are desirable. Of our datasets, this only occurs in arthropod COI, because no secondary ranks are used in the fungal ITS data. Second, there may be genuine uncertainty about a species’ relationship to other organisms, such that taxonomists have chosen not to classify at all ranks. This situation is more common in our fungal ITS dataset. While the taxonomic annotations could contain some errors, for the purposes of this evaluation we treat them as the ground truth. To reflect realistic differences between training and test sets, we used sequences from different countries for COI and sequences derived from different methodological approaches for ITS.

#### COI sequences: FinBOL and GBOL

For COI, we used DNA sequences from the Finnish Barcode of Life (FinBOL; Roslin et al., 2022) as the training set and from the German Barcode of Life (GBOL; German Barcode of Life Consortium, 2011) as the test set. FinBOL and GBOL aim to create comprehensive DNA barcode reference libraries for multicellular organisms in Finland and Germany, respectively, using targeted sequencing of expert-identified specimens. Because they focus on separate regional faunas and, to some degree, reflect the research priorities of their respective taxonomic communities, the sets have only a partial taxonomic overlap.

For FinBOL, sequences and taxonomic annotations (including ranks class, order, family, subfamily, tribe, genus, and species) were retrieved from the BOLD (Ratnasingham and Hebert, 2007; Ratnasingham et al., 2024) data package 29-Mar-2024 (BOLD Systems, 2024) using the project code ‘DS-FINPRO’. For GBOL, sequences were retrieved from the GBOL web portal (German Barcode of Life Consortium, 2011) using the filters “Sub-/Phylum: Arthropoda”, “Collected: 11”, and “With Barcode: 11” to retrieve COI barcodes from a maximum of eleven specimens per species within Arthropoda. To avoid discrepancies between the taxonomy of the training and test sets, the Process-ID corresponding to records in BOLD was used to retrieve taxonomic information for each sequence in the test set.

Secondary taxonomic ranks, here subfamily and tribe, are not assigned to all taxa in the arthropod taxonomy. To represent taxa without secondary ranks, we used *dummy* as placeholder taxa. In our evaluation of COI classifications, we treated these placeholder taxa as equivalent: for example, the genera *Acanthosoma* and *Elasmostethus* were classified in the subfamily *Acanthosomatinae*, but did not have a tribe classification. At the tribe rank, they were therefore both represented as `Acanthosomatinae_dummy_tribe`. We avoided the use of the more standard taxonomic term *“-incertae sedis”*, meaning “of uncertain placement”, because the MycoAI classifier ignores taxa which end with this term, leading to unexpected poor performance in our tests at these ranks.

Several classifiers we tested, including dnabarcoder, SINTAX, and MycoAI, require explicitly named primary Linnean ranks. For these algorithms, we recoded the ranks above genus level as tribe → family, subfamily → order, family → class, order → phylum, and class → kingdom.

For both training and test sets, we retained only records that 1) were identified to the species level; 2) did not belong to a “placeholder” taxon at the species level; 3) had been annotated as being a sequence of the COI-5P barcode region; and 4) were at least 600 bp in length. Training and test sequences were aligned to the Folmer barcode region (Folmer et al., 1994) and amino acid translations were generated using MACSE v2.07 (Ranwez et al., 2018) and the COI coding sequence from *Drosophila melanogaster* (NCBI accession NC_024511.2, positions 1474..3009) as a reference. Sequences with less than 600 bases in the nucleotide alignment after removal of gap columns were removed from both nucleotide and amino acid alignments, resulting in 35 603 training and 26 925 test sequences; see Supporting Information S1 for further details. The final COI test set included novel taxa at all ranks from class to species, as displayed in Fig. 1.

**Figure 1:**
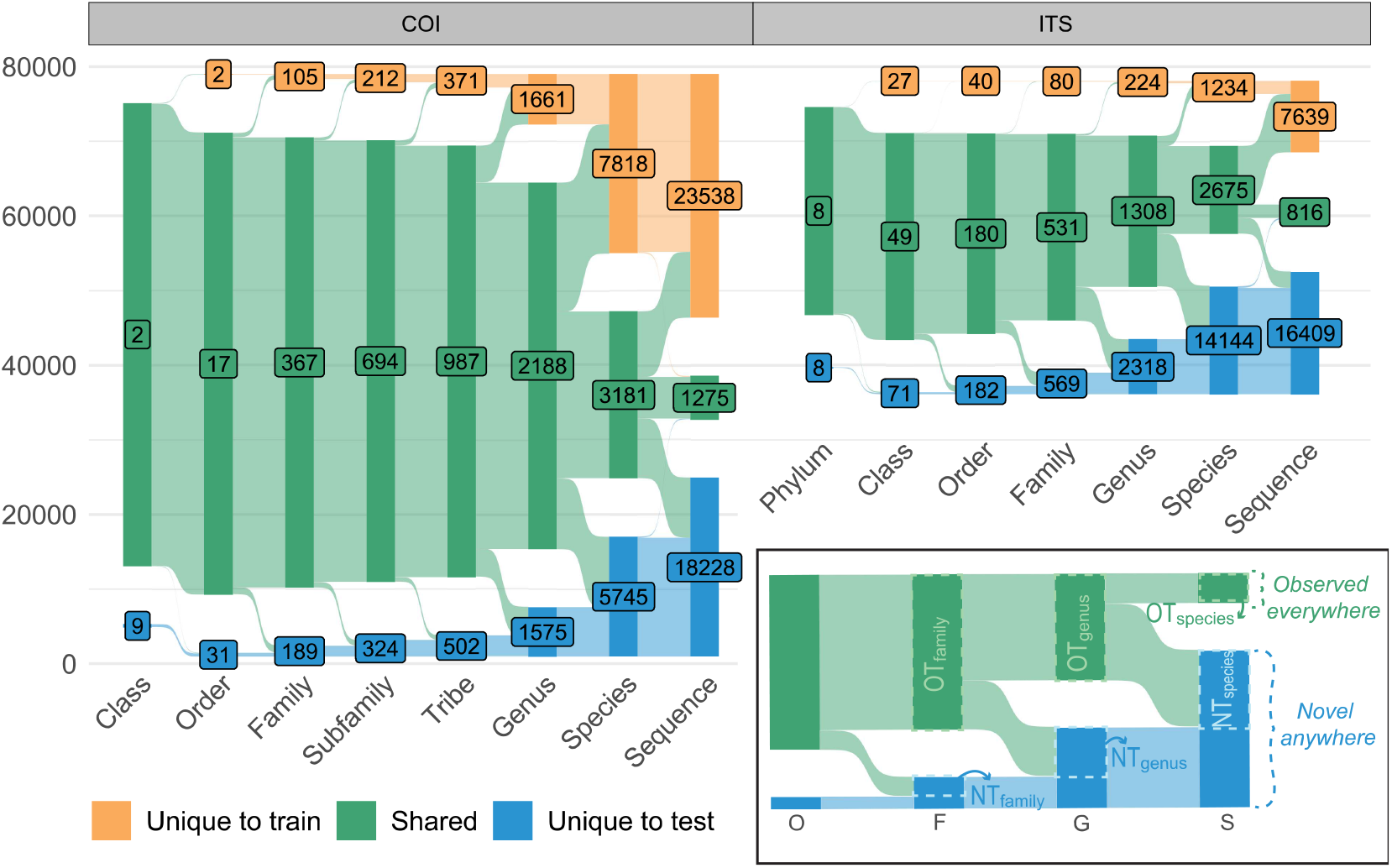
Taxonomic overlap between training set and test set for COI and ITS sequences. The x-axis displays the taxonomic ranks, with an additional column for unique DNA sequence variants. The height of columns and ribbons correlates to the number of sequences belonging to either shared taxa (green), taxa unique to the training set (yellow) or taxa unique to the test set (blue). Numbers show the count of unique taxa (or, for the last column, sequences) at each rank and group. The inset demonstrates how we partitioned the test data into *observed everywhere* and *novel anywhere*, as well as *observed taxa* (OT) and *novel taxa* (NT) at each rank, using ITS test sequences across ranks order to species as an example.

We generated the testshort data set for COI by replacing the first 240 positions of the nucleotide test alignment and the first 80 positions of the amino acid test alignment (i.e., up to and including the BF3 primer binding site; Elbrecht et al., 2019) with gaps (“-”). Trimmed but unaligned training, test, and testshort sequences were generated by removing all gap characters from the aligned sequences.

#### ITS sequences: Westerdijk culture collection and Unite reference sequences

For ITS, we used DNA sequences from the Westerdijk Fungal Biodiversity Institute culture collection of yeasts (Vu et al., 2016) and filamentous fungi (Vu et al., 2019) as the training set. Our test set consisted of curated *reference sequences* from the UNITE database (Abarenkov et al., 2023). These are sequences that have been selected manually to represent species hypotheses. The Westerdijk Institute maintains the largest culture collection of living fungi in the world, while the UNITE database is a community-curated database of eukaryotic ITS sequences generated from diverse substrates and methods. The implied differences in scope and methodological breadth result in a partial taxonomic overlap.

For the training set, sequences were retrieved from NCBI Bioprojects PRJNA351778 (https://www.ncbi.nlm.nih.gov/bioproject/351778) and PRJNA422523 (https://www.ncbi.nlm.nih.gov/bioproject/422523) and matched with taxonomic annotations (including ranks kingdom, phylum, class, order, family, genus, and species) from the UNITE database (Abarenkov et al., 2023). For the test set, we downloaded the UNITE 10.0 general FASTA release from https://doi.org/10.15156/BIO/2959332 and selected sequences annotated as reference sequences (“RefS”). Many sequences from the Westerdijk culture collection also occur as reference sequences in UNITE. To avoid inclusion of the same record in the train and test sets, we removed duplicate records from the training set if multiple sequences were available for that species hypothesis. If the duplicate sequence was the unique representative of a species hypothesis, we instead removed it from the test set.

We obtained taxonomic information for Unite species hypotheses from Abarenkov (2024). For fungal species which do not include all primary taxonomic ranks in their classification, the Unite database uses *incertae sedis* names as placeholder taxa, paired with the name of the closest enclosing taxon and its rank. For example, the genus *Ceratocladium* is classified in phylum *Ascomycota*, but not in any of the many described classes, orders, or families within *Ascomycota*, so its family is listed in Unite as `Ascomycota_fam_Incertae_sedis`. Multiple taxa with the same placeholder cannot be assumed equivalent; for instance, genus *Septogloeum* is also listed at the family level as `Ascomycota_fam_Incertae_sedis`, but this does not indicate that the genera *Ceratocladium* and *Septogloeum* belong to the same natural group at the family level. To disambiguate such cases, we appended the nearest named lower rank to each *Incertae sedis* label, in the example case creating the family-level placeholders `Ascomycota_fam_Incertae_sedis_Ceratocladium` and `Ascomycota_fam_Incertae_sedis_Septogloeum`.

For both datasets, we extracted the ITS region with ITSx (Bengtsson-Palme et al., 2013), which uses profile hidden Markov models (HMMs) to detect the flanking SSU and LSU regions. When the included fragments of SSU or LSU are too short, ITSx sometimes fails to detect them. To solve the issue, we used cutadapt (Martin, 2011) to trim sequences matching the end of SSU, and LSUx (Furneaux et al., 2021) to detect and remove the beginning of LSU. Finally, we created testshort sequences including only ITS2 by deleting all nucleotide bases up to the end of 5.8S, as detected by Rfam covariance model RF00002 (Kalvari et al., 2021).

### 2.2 Classification algorithms

We trained each classification algorithm on the training set and then classified all sequences in the test and testshort sets, as well as the amino acid versions of test and testshort for COL Some methods feature many adjustable parameters for training and/ or classification. To ensure a fair comparison, we used default settings when possible, or parameter values that we believe reflect standard usage. Hence, we did not tune parameters to optimize results on our particular dataset, nor did we modify the algorithms from their most recent publicly available versions to improve their performance. Moreover, for algorithms that produce multiple alternative prediction per query at each rank, we selected the prediction with the highest probability or confidence score, while remaining consistent with the predictions at higher ranks. We determined whether each prediction was correct by comparing it with the taxonomic annotation in the original test set. If the true taxon was unobserved in training, we considered the classification as correct if the taxon was predicted novel at the correct rank and all higher ranks were predicted correctly.

Some algorithms do not predict novelty but can refrain from making predictions for sequences with high classification uncertainty. Since a high proportion of missing predictions can inflate classification accuracy, we distinguish between results that allow missing predictions (MP) and those where predictions are enforced for all query sequences (FP). In the MP setting, we applied confidence thresholds for classification algorithms as recommended in the original publication, and interpreted classifications below this threshold, as well as missing predictions, as neither correct nor incorrect. In the FP setting, we did not use any confidence threshold, and interpreted missing predictions as the algorithm classifying the sequences as a novel taxon.

Algorithms were executed on a compute node of a Linux high-performance computing cluster equipped with two Intel Xeon Gold 6230 CPUs for a total of 2×20 processing cores at 2.1 GHz. For each method, the full training and classification task was run with allocations of 1, 4, 16, and 40 processor cores to test parallel scalability. If GPU execution was supported, methods were additionally run in allocations with 1 CPU core and 1 Nvidia V100 GPU. We adjusted the memory and time allocations as needed for the algorithms to complete, with a maximum limit of 192 GiB of memory and 14 days of processing time. Elapsed time, maximum memory usage, and average processor usage of the training and classification steps were measured using the Gnu time utility. Disk space required by the trained model was also recorded. See Table 1 for an overview of the algorithms we tested, and Supporting Information S1.2 for execution details for each algorithm. Scripts used to install, train, and test each algorithm are publicly available at the following GitHub repository.

#### 2.2.1 Leveraging additional information

PROTAX and EPA-ng present opportunities for the user to augment the model with additional information. For PROTAX, we evaluated a base case, trained only on the training sequences, as well as an augmented case, where we supplied the full arthropod taxonomy from BOLD snapshot 29-Mar-2024 (Ratnasingham et al., 2024; BOLD Systems, 2024) for the COI case, and the full fungal taxonomy from the Unite SH training data (Abarenkov, 2024) for the ITS case. For EPA-ng, we tested three different reference trees: one generated entirely de-novo from the reference database with no constraints, one built with full taxonomic constraints, and one built with constraints based on previously established phylogenies for which relationships between families are resolved. We refer to the first case as the *free tree*, the second case as the *taxonomically constrained tree*, and the third case as the *phylogenetically constrained tree*. Phylogenetic placement algorithms typically require that the query and reference sequences are globally aligned, making them unsuitable for use with barcode regions for which multiple sequence alignment is not feasible, such as ITS. Therefore, we only tested this class of models for COI.

### 2.3 Data partitions

To explore the differences in algorithm performance on query sequences belonging to either observed or novel taxa, we split sequences into different partitions. First, the *observed everywhere* partition included test sequences from taxa represented in the training data at all taxonomic ranks, while the *novel anywhere* partition included sequences from taxa that were novel on one or more ranks (Fig. 1). Second, to explore separately how well algorithms predicted each taxonomic rank, we split sequences into the *observed taxa* and the *novel taxa* partitions. *Observed taxa* included, for each rank, all test sequences belonging to an observed taxon on that particular rank. For example, at order level, it included test sequences belonging to orders represented in the training data, regardless of whether that sequence was from an observed taxon at lower taxonomic ranks or not. Similarly, *novel taxa* included all test sequences at each rank that belonged to a taxon that was novel at that particular rank, but not at any higher rank. For instance, each sequence in *novel taxa* at the family level belonged to observed orders but novel families. Consequently, the total number of sequences in the *observed taxa* and *novel taxa* partitions varied depending on the rank under consideration.

### 2.4 Performance metrics

To differentiate between algorithm performance, we calculated a classification accuracy where correct novelty prediction was labeled as a true positive. In other words, this metric was defined as

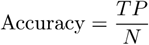

where *TP* was the number of correct predictions, including novel sequences correctly predicted as novel, and *N* was the number of sequences in the test set. We also calculated three further metrics of performance: 1) the *over-classification rate*, where taxonomic names are assigned to novel taxa; 2) the *under-classification rate*, where observed taxa are predicted as novel; and 3) the *misclassification rate*, where incorrect taxonomic names are assigned to observed taxa, using the definitions by Edgar (2018). Error rates are evaluated under the MP setting, using recommended confidence thresholds when present. For observed and novel taxa, we further calculated marginal and conditional recall. Recall is a performance metric used to measure of a model’s ability to identify positive instances out of all true instances in the dataset (also known as sensitivity). Here, we defined *marginal recall* for each taxonomic rank as

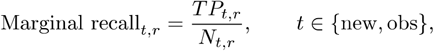

where *TP*_new,*r*_, *TP*_obs,*r*_ were the number of correct taxonomic predictions of sequences belonging to the groups of novel and observed taxa, respectively, at rank *r*, and *N*_new,*r*_, *N*_obs,*r*_ were the number of sequences in the test set belonging to the groups of novel and observed taxa at rank *r*, respectively. Note that *TP*_new,*r*_ +*TP*_obs,*r*_ = *TP* and *N*_new,*r*_ *+ N*_obs,*r*_ = *N* at any rank. For *conditional recall*, we considered only cases where the predictions for the higher ranks were correct. That is,

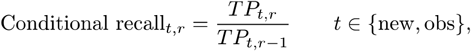

where *TP*_*t,r*−1_ is the number of sequences correctly predicted at rank *r* − 1 for group *t*.

We also calculated prediction coverage, defined as the proportion of sequences for which an algorithm assigned a taxonomic label, across all possible confidence thresholds. For algorithms that did not produce confidence estimates, coverage was calculated as a single value. In both cases, we considered the MP setting, thus allowing missing predictions also in the calculation of the coverage. However, for algorithms with a recommended confidence threshold to accept predictions, we computed prediction coverage across the full range of confidence thresholds.

Finally, we evaluated the calibration of the confidence estimated by plotting the cumulative probability of the predictions against the cumulative proportion of cases where the prediction was correct. For well-calibrated algorithms, probability and proportion of correct predictions are equivalent: in cases where the prediction probability is 90%, the predictions are correct 90% of the time. Graphically, this results in a line close to 45 degrees. Instead, deviations from the 45-degree line are interpreted as under- or over-confidence. For algorithms missing confidence estimates, we assigned 100% confidence to all predictions.

## 3 Results

### 3.1 Classification accuracy and recall

When classifying the *observed everywhere* partition under the FP setting, where we enforced predictions for all sequences, most algorithms exceeded 93% and 78% accuracy at the genus and species level, respectively, for both COI and ITS sequences (Fig. 2a, for MP setting see Fig. S2). For the *novel anywhere* partition, all algorithms exhibited a lower classification accuracy than for *observed everywhere*, ranging between 9-58% accuracy at the genus level for COI, and 12-40% for ITS. Models that explicitly distinguished between classification uncertainty and novel taxon predictions, i.e., EPA-ng (COI only), BayesANT, and PROTAX, classified *novel anywhere* with accuracies between 12-44% for COI, and 0-29% for ITS. For COI, EPA-ng consistently achieved the highest classification accuracy across ranks, a result that held for both the taxonomically and the phylogenetically constrained reference tree. Phylogenetic placement on the free tree resulted in slightly lower accuracies, but it was still among the top performing algorithms. For ITS, composition-based classifiers exhibited the highest prediction accuracy for *novel anywhere*, with nearly identical accuracies for IDTAXA, RDP-NBC, and SINTAX (Fig. 2a).

**Figure 2:**
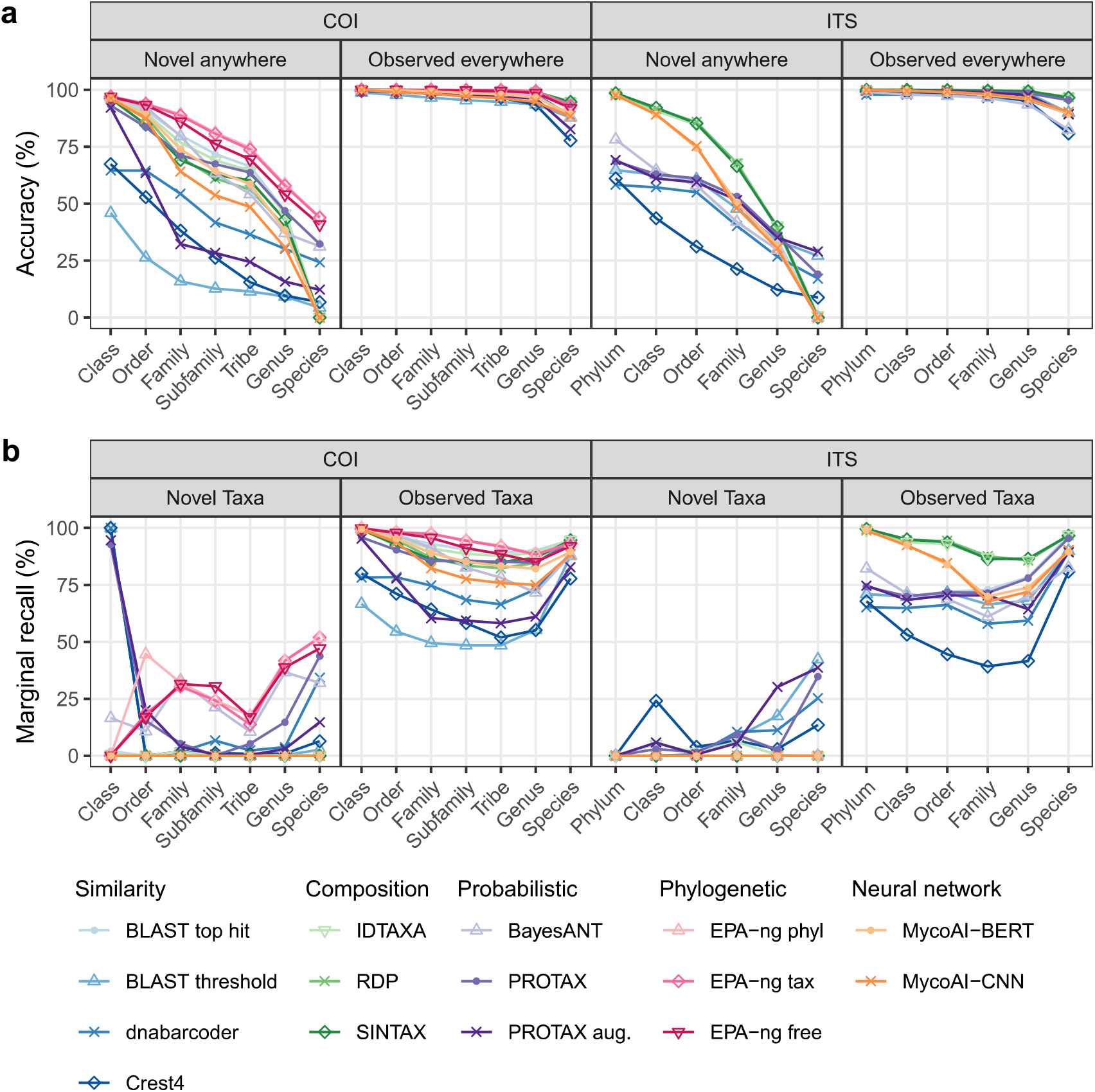
Accuracy and marginal recall of classification algorithms across taxonomic ranks under the FP setting, where predictions were enforced for all query sequences. **a** shows accuracy, defined as the proportion of all taxonomic classifications that were correct for observed (right) and novel (left) species. **b** shows marginal recall, calculated as the proportion of correct predictions relative to the total number of sequences belonging to either observed (right) or novel (left) taxa at that rank.

The *observed taxa* partition contained sequences which may be novel at lower ranks and classification of these sequences may be more difficult than those observed at species level. Accordingly, all algorithms showed lower marginal recall than accuracy at the corresponding rank (Fig. 2b). This suggests that higher-level classifications were more error-prone when the sequence belonged to a novel taxon at a lower rank, compared to when a species-level match was available. Compared to the *observed everywhere* partition, the difference between algorithms was substantially larger for *observed taxa*. In general, similarity-based classifiers exhibited the lowest marginal recall for *observed taxa* for both COI and ITS, except BLAST top hit, which was consistently among the top-performing algorithms across all ranks for COI, and at ranks below order for ITS. For *novel taxa*, only EPA-ng, BayesANT, and PROTAX explicitly predicted taxonomic novelty. Some similarity-based algorithms can abstain from making predictions, which we interpreted as novelty in the FP setting. Consequently, as expected, all composition-based and neural network classifiers exhibited zero marginal recall for *novel taxa*. Among the algorithms that explicitly predicted novelty, EPA-ng and PROTAX exhibited the highest marginal recall for COI and ITS, respectively (Fig. 2b). BayesANT achieved markedly better performance for COI sequences than ITS sequences, where it failed to recover any novel taxa. For COI, BayesANT exhibited the highest conditional recall across ranks from family to species, suggesting it successfully predicted novelty in most cases where the higher ranks were predicted correctly (Fig. S3). For COI, EPA-ng, BayesANT, and PROTAX all overestimated the number of novel taxa at the family level and below (Fig. S4). For ITS, PROTAX overestimated and BayesANT underestimated the number of novel taxa across all ranks.

When evaluated on the testshort dataset, designed to mimic shorter metabarcoding reads, most algorithms classified *observed everywhere* with accuracies similar to full-length reads, with the exceptions of MycoAI for both markers and DNABarcoder and RDP-NBC for ITS (Fig. S5). Notably, DNABarcoder assigned almost all ITS testshort sequences from *observed everywhere* to a novel class, yielding accuracy close to zero. For *novel anywhere*, classification accuracy declined on short reads compared to full-length reads, particularly for ITS at higher taxonomic ranks.

Translating COI to amino acid sequences yielded lower classification accuracies of *observed everywhere* (Fig. S6). However, BayesANT and Crest4 exhibited higher accuracies compared to nucleotide sequences for *novel anywhere*, especially at higher ranks.

### 3.2 Error rates

Most models showed very low misclassification rates (<5%, for COI and <14% for ITS; Fig. 3). Higher error rates were observed for MycoAI and BLAST top hit across both markers, and for BayesANT on ITS. In addition, at the ITS species level, RDP-NBC and SINTAX showed elevated error rates, misclassifying 20% and 14% of *observed everywhere*, respectively. Overall, over-classification and under-classification showed opposing patterns, highlighting the trade-off between the two. Even under the MP setting, there is no recommended confidence threshold or stopping rule for MycoAI or BLAST top hit, thus resulting in 100% over-classification for MycoAI for both markers, and nearly 100% over-classification for BLAST top hit and COI. For ITS, BLAST top hit had a higher proportion of missing predictions, resulting in lower over-classification rates. Among the algorithms that included a stopping rule, EPA-ng had the highest over-classification rate for COI. In contrast, BayesANT achieved both a low over-classification rate and a relatively low under-classification rate for COI. For ITS however, BayesANT over-classified most novel taxa. RDP-NBC and SINTAX had overall low over-classification rates for COI, except at the class level, where they over-classified 73% and 98% of novel taxa, respectively.

**Figure 3:**
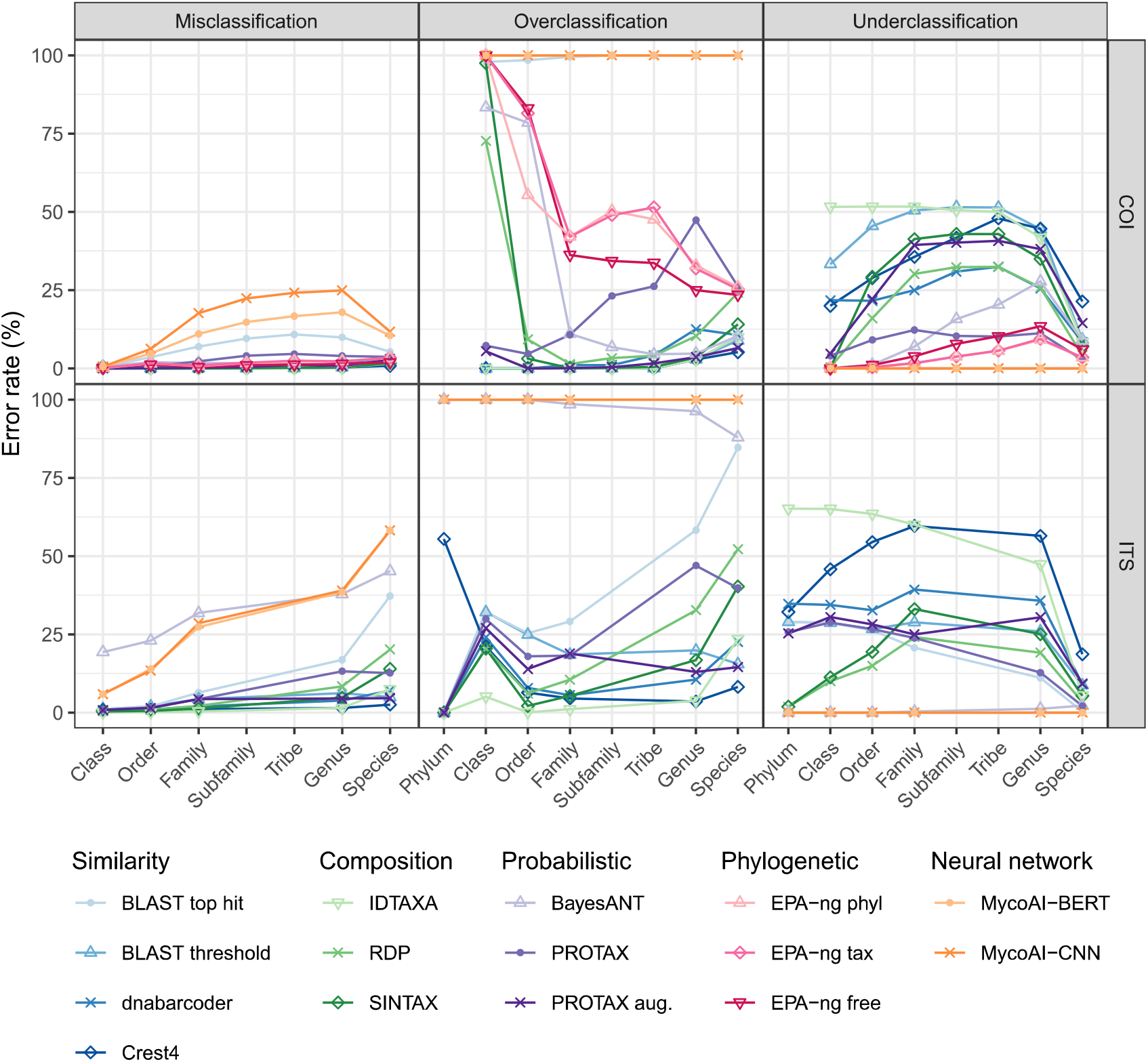
Rates of mis-, over-, and under-classification of COI and ITS sequences across classification algorithms and taxonomic ranks. Error rates are shown under the MP setting, allowing missing predictions and implementing confidence thresholds for algorithms that recommend them.

### 3.3 Calibration of prediction probabilities

For algorithms that produced prediction probability estimates, probabilities were well calibrated for the *observed everywhere* partition, with prediction probability and prediction accuracy following a path close to the 45-degree line (Fig. 4). For the *novel anywhere* partition, MycoAI produced well-calibrated probability estimates for both COI and ITS predictions. The remaining algorithms exhibited either excessive confidence or a lack of it. In general, algorithms were more over-confident for ITS than for COI, especially at lower ranks (Fig. S7-S8).

**Figure 4:**
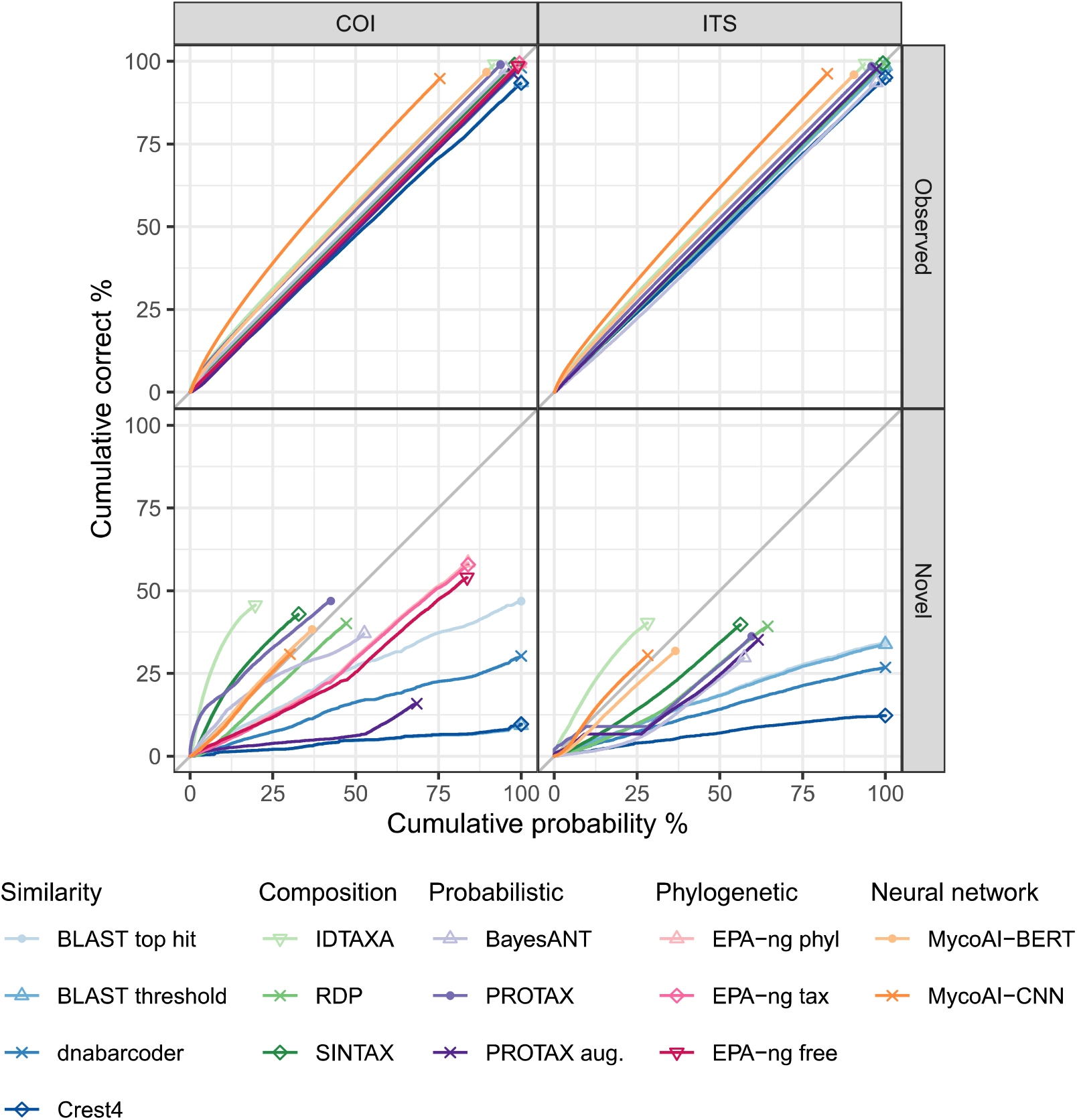
Calibration of different classification algorithms at genus-level taxonomic predictions (for other ranks, see Fig. S7 and S8). The x-axis shows the cumulative probability of the prediction with the highest probability or confidence score and the y-axis shows the cumulative proportion of correct predictions. Models are well calibrated if the line follows the identity line, here shown in gray. The y coordinate of the point at the end of the line displays the average accuracy. Calibration is shown for COI and ITS sequences, and for the partitions *observed everywhere* and *novel anywhere*. Calibration results are under the FP setting, i.e. where predictions are enforced for all query sequences.

### 3.4 Classification coverage

There was a clear trade-off between accuracy and prediction coverage, where increasing confidence thresholds resulted in lower coverage but higher classification accuracy, as displayed in Fig. 5. The exact relationship between prediction coverage and accuracy varied between algorithms; some achieved moderate accuracies at high coverage, while others required aggressive filtering to achieve the same accuracy. Differences were especially pronounced for COI at the species level, where EPA-ng preserved high accuracy even at 100% prediction coverage. Algorithms without confidence estimates, such as BLAST top hit, appeared as fixed points in this trade-off space, classifying relatively few sequences but often with high correctness. Importantly, there was no consistent threshold that yielded a high degree of correct classifications across algorithms or genetic markers. The optimal operating point instead depends on whether accuracy or recall is more important for the application.

**Figure 5:**
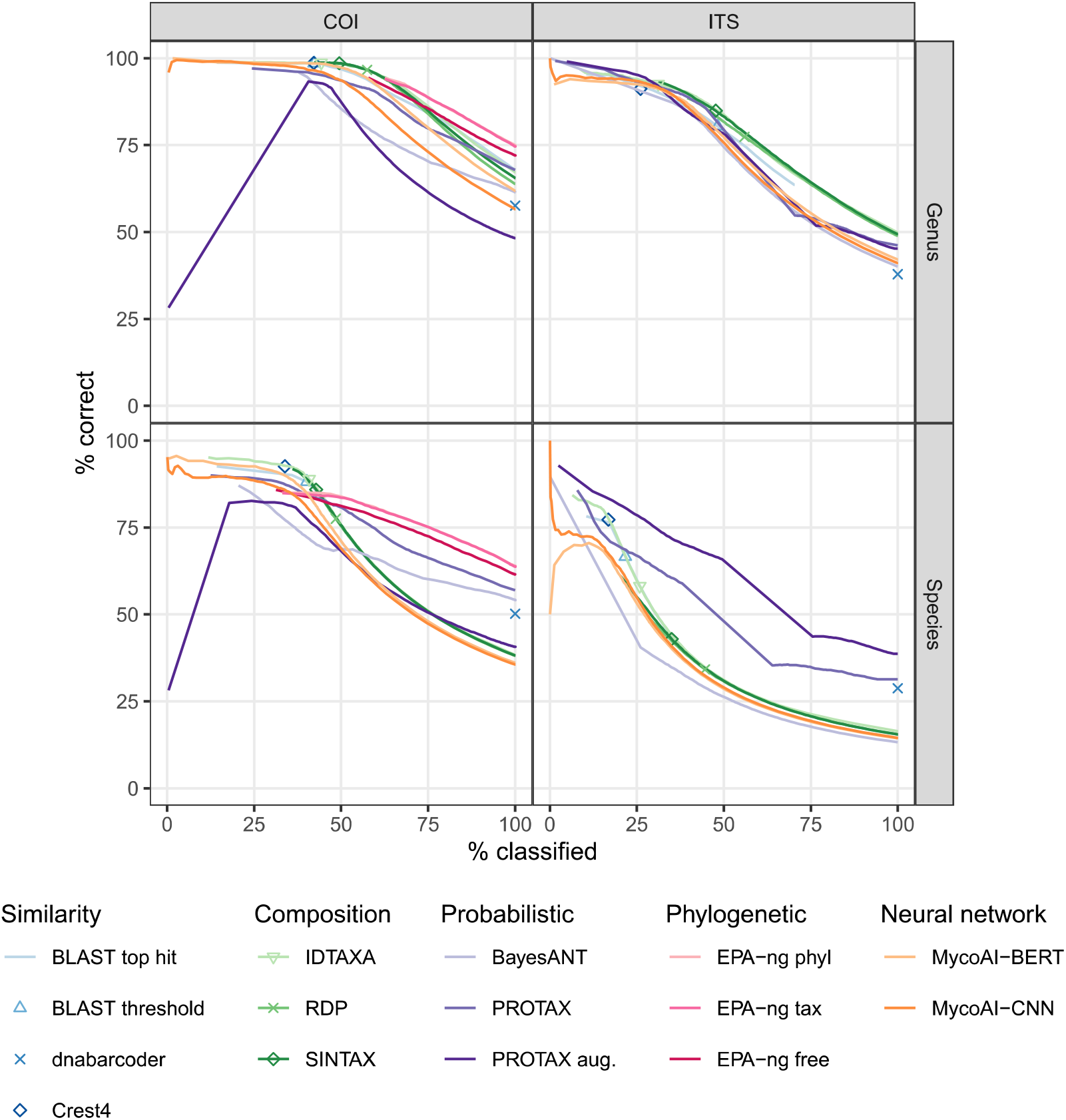
Classification accuracy as a function of prediction coverage for different algorithms. The x-axis shows the proportion of sequences classified and the y-axis shows the proportion of correct predictions. Continuous lines indicate algorithms with probability or confidence estimates, where varying the threshold yields a trade-off between coverage and accuracy. Single points represent algorithms without confidence estimates, corresponding to a fixed trade-off. We allowed missing predictions, but did not implement confidence thresholds for algorithms that recommend them. However, recommended thresholds are indicated by points along the continuous line.

### 3.5 Computational resources

There was no clear relationship between classification performance and computational resource use. We report all metrics in Table S3 in the Supporting Information. Lightweight methods such as SINTAX achieved high accuracy with minimal runtimes and were the fastest overall when accounting for both training and classification, while resource-intensive algorithms such as MycoAI required substantially more time and memory for training without yielding superior accuracy. Once trained however, classification with MycoAI was quick, with the CNN model having the shortest classification runtime overall. EPA-ng was substantially slower on amino acid sequences than nucleotide sequences, due to the increased complexity of the substitution model. Most other algorithms were faster on translated sequences, likely due to their reduced length. Access to multiple CPU cores or a GPU reduced runtimes for most algorithms, especially during the classification phase, whereas fewer algorithms benefited substantially from parallelization during initialization or training, with MycoAI being a notable exception. Since training is typically a one-time cost, scalability during classification may be the more critical factor for real-world applications. Model size on disk also varied widely, with MycoAI and BayesANT models occupying 0.2-0.4 GB compared to less than a hundred MB for many similarity- or composition-based classifiers, which may make the latter more practical for deployment in large-scale or resource-limited settings.

## 4 Discussion

DNA barcoding has proved a versatile tool for biodiversity research, and its applications are continuously expanding and developing following advances in sequencing technologies (Gostel and Kress, 2022; Van Klink et al., 2022; Niskanen et al., 2023). For many purposes, however, reliable taxonomic assignment of the resulting barcode sequences is paramount for downstream analyses. The primary factor determining classification accuracy is the completeness of the reference database, as emphasized in previous studies (Sickel et al., 2015; Taberlet et al., 2012; Edgar, 2018) and strongly supported by our result in Fig. 2. However, we have demonstrated that the sensitivity to incomplete reference data varies between taxonomic classifiers and that differences in performance between algorithms are more pronounced when species-level matches are lacking. Our results highlight that taxonomic classifiers vary in how they balance different types of classification errors, emphasizing the importance of careful selection of algorithms suitable for both the genetic marker targeted and the research question at hand.

### 4.1 High classification accuracy comes at the cost of prediction coverage

We found a general *trade-off* between classification accuracy and prediction coverage across all methods (Fig. 5). This outcome is intuitive: algorithms that abstain from making predictions when uncertain have a lower risk of misclassification. When dealing with novel taxa, this corresponds to the balance between under- and over-classification. Under-classification increases accuracy at higher ranks by withholding predictions when uncertainty is high, while over-classification increases coverage but also increases the chance of errors at lower ranks. Because algorithms vary in how they navigate this trade-off, their suitability depends on the objectives of a given study. For instance, biodiversity monitoring or community surveys may benefit from a high prediction coverage at the expense of accuracy, as detecting rare taxa is crucial (Mouillot et al., 2013; Soliveres et al., 2016). Conversely, applications such as network analysis may prioritize accuracy, since misclassifications can create spurious links between taxa (Cuff et al., 2022). To support informed decision-making, we argue that classification algorithms should provide well-calibrated probabilities or confidence estimates.

Clearly reporting classification uncertainties not only enhances scientific reliability but also enables researchers to incorporate these probabilities into downstream analyses, thereby refining subsequent inferences. When reference databases are incomplete, algorithms should additionally distinguish between novelty and classification uncertainty, improving transparency and interpretability. Algorithms that rely on a single threshold to accept or reject classification must optimize thresholds to balance between different error types, often resulting in conservative cut-offs as observed in Fig. 5. Treating confidence scores below such thresholds as indicative of novelty can thus inflate estimates of unseen taxa. For instance, both BLAST threshold and dnabarcoder largely over-assigned DNA sequences to novel branches (Fig. S4), highlighting that many “under-classifications” reflect the stringency of the applied thresholds rather than true novelty. On the other hand, algorithms that explicitly account for the possibility of novel taxa and distinguish between novelty and classification uncertainty greatly enhance interpretability, enabling researchers to disentangle true biological signal from methodological conservatism and thereby draw more robust ecological and evolutionary inferences. Consequently, researchers should favor classification approaches that explicitly handle novelty when working in geographic regions or with taxonomic groups that are known to be underrepresented in reference databases.

### 4.2 Different algorithm classes perform best on different genetic markers

Genetic markers vary in their suitability for taxonomic classification of different taxonomic groups, owing to lineage-specific rates of evolution and variation in sequence composition (Hebert et al., 2003; Pentinsaari et al., 2016; Schoch et al., 2012). We evaluated classification algorithms on two widely used DNA barcodes chosen specifically for their contrasting characteristics: the COI barcode as a protein-coding gene with strong evolutionary constraints on the amino acid sequence and length, and ITS as a non-coding region of variable base composition and length.

While every method reached a high classification accuracy for observed species (i.e. the *observed everywhere* partition), we detected substantial differences in performance across genetic markers for novel taxa. For ITS, composition-based algorithms achieved the best results (e.g., Fig. 2, green colors). This is likely due to the high sequence variability and the resulting difficulty in establishing position homology for distant species, which prevents reliable sequence alignments. Hence, *k*-mer-based approaches likely benefit from the strong signal provided by recurring short motifs, which are more indicative of relatedness in a highly variable marker like ITS. Instead, the evolutionary constraints in COI may produce short sequence matches by pure chance. In this regard, the best performance for COI was achieved by phylogenetic placement (Fig. 2, pink colors), which explicitly employs evolutionary models to infer taxonomic relationships. We did not evaluate this class of algorithms for ITS due to the difficulties in providing a global alignment. However, alternative phylogenetic placement algorithms relying on *k*-mer composition are available (Balaban et al., 2019), and could be an interesting future avenue to explore for taxonomic classification of ITS sequences. Overall, our result indicates that a better understanding of both sequence composition and evolutionary constraints can help researchers select the most appropriate algorithm for their specific application.

### 4.3 Classification performance is not determined by computational resource use

We did not observe a general improvement in classification performance at the expense of higher computational costs. Indeed, the best performing algorithms for ITS included SINTAX, which had the lowest combined training and classification time, whereas computationally intensive phylogenetic placements were particularly rewarding for COI classifications. To save computational resources, large COI datasets may therefore benefit from a two-step strategy: first applying a fast algorithm such as SINTAX with a high confidence threshold to classify observed species, and then using the more costly EPA-ng to classify only those sequences that could not be confidently assigned, as in Sundh et al. (2025). Because SINTAX confidence estimates for COI were consistently under-confident (Fig. 4), a high confidence threshold would yield very few mis- or over-classifications.

Although we assessed scalability in terms of available computational resources, we did not specifically examine how the models scale with the size of the reference database, which is an important practical consideration. Realistically, the size of the present databases might limit the use of resource-heavy algorithms. For phylogenetic placement, for example, it may be necessary to preselect representative taxa for inclusion in the reference tree to make it computationally tractable.

### 4.4 Limited impact of read length on classification performance

In many applications, shorter sub-regions of the DNA barcode are employed. Shorter reads often amplify more reliably than full-length barcodes (Meusnier et al., 2008) and are therefore preferable when working with degraded DNA. They can also be sequenced more cost-effectively on high-throughput platforms, making them attractive for large-scale barcoding studies. For most algorithms, we observed minimal differences in classification performance between short reads and full-length barcodes, suggesting that the regions targeted by the shorter reads captured most of the variation relevant for species classification. Previous studies have reported similar results for both COI (Yeo et al., 2020) and ITS (Badotti et al., 2017).

Notably, dnabarcoder failed to classify 99% of ITS short reads as belonging to the kingdom Fungi, resulting in near-zero accuracy, and MycoAI showed significantly reduced performance on short reads across genetic markers and data partitions. However, we trained all classifiers on the full-length barcodes and did not retrain them prior to classifying the short reads. The observed performance drop may therefore reflect sensitivity to mismatches between training and query sequence length, rather than an inherent inability of these algorithms to annotate short sequences.

### 4.5 Barcode-based classifications under taxonomic uncertainty

For successful taxonomic classification of novel taxa from DNA barcodes, the genetic variation of the barcode must reflect the relatedness of taxonomic groups. For observed species, a genetic marker is generally suitable for taxonomic classification if the interspecific variation exceeds the intraspecific variation. For novel taxa, however, this criterion must be fulfilled also at higher taxonomic levels, so that the barcode sequence composition is sufficient to determine the correct genus, family, or even higher rank without an exact species match. This task is further complicated by the absence of universal divergence thresholds that reliably indicate the lowest shared taxonomic rank between two taxa (Edgar, 2018). In other words, even with the same genetic marker, a 3% divergence may correspond to taxa that share either a genus or a family. Additionally, there are incongruencies between currently accepted Linnean taxonomic classifications and molecular phylogenies, and taxonomic relationships are continually revised as new phylogenetic evidence corrects previous classifications (e.g., Möckel et al., 2022; Chen et al., 2021; Johnston et al., 2024; Ji et al., 2023). Beyond inconsistencies between reference databases, which differ in how frequently they update their taxonomies, such incongruencies can undermine the consistency of barcode-based classification by violating the fundamental assumption that taxonomy accurately reflects evolutionary relatedness. These challenges highlight the appeal of methods that infer evolutionary relationships directly rather than relying solely on taxonomic labels. Phylogenetic placement is one such approach, integrating evolutionary signal into the classification process. However, its utility for taxonomic classification ultimately depends on how closely the reference taxonomy reflects true phylogenetic relationships.

Phylogenetic placement accuracy may also depend on the quality of the reference tree. Previous studies have shown that phylogenetic backbones can improve tree inference from DNA barcodes by supplementing their limited signal for deep evolutionary relationships (Talavera et al., 2022; **Liu** et al., 2019). To examine whether a more accurate phylogenetic representation of higher-level relationships translates into improved classification performance, we evaluated EPA-ng using two reference trees: one constrained by taxonomy alone and one incorporating phylogenetic relationships on the family level and above. Hope-inspiringly, we observed minimal differences in classification results between the two reference trees, suggesting that constraining the tree to match taxonomy is generally sufficient for accurate classifications.

### 4.6 Recommendations for algorithm selection

Our results clearly show that the choice of taxonomic classifier is less critical when reference libraries are complete, allowing researchers to prioritize computational efficiency. Nevertheless, even with comprehensive reference data, it remains important to choose an algorithm that provides confidence estimates to flag uncertain cases. For this purpose, our results highlight SINTAX as a fast, memory-efficient option that produces well-calibrated, albeit slightly conservative, classification confidence estimates.

When reference libraries are incomplete, our results suggest that phylogenetic placement methods (e.g., EPA-ng) are preferable for COI barcodes, whereas composition-based methods (e.g., RDP-NBC, SINTAX, and IDTAXA) perform best for ITS barcodes. While our findings are specific to COI and ITS, the observed trends might be indicative for other barcodes sharing similar characteristics, though this remains to be validated. Under incomplete reference libraries, we further highlight the importance of selecting algorithms that explicitly distinguish between classification uncertainty and novel taxa, such as EPA-ng, PROTAX, and BayesANT. However, for ITS, no algorithm consistently achieved reliable identification of novel taxa across ranks. Consequently, the interpretation of whether uncertain classifications indicate true novelty is left to the user, underscoring the importance of transparent criteria for assigning novelty and clear reporting of well-calibrated confidence estimates.

To reduce computational demands for large COI barcode datasets, we recommend a two-step strategy, as also suggested by Sundh et al. (2025). First, apply a fast and resource-efficient method such as SINTAX with a high confidence threshold to classify most observed species. Then, apply phylogenetic placement to the remaining uncertain cases to optimize classification. Although not evaluated here, the computational efficiency of phylogenetic placement may improve if the reference tree is built from a preselected set of representative sequences, that is, one sequence per species, rather than the full set used in this study.

Finally, our evaluations considered off-the-shelf approaches to taxonomic classification that leveraged default settings. In principle, one can further improve performance by properly tuning model-specific parameters to the training data and the specific marker, noting that some algorithms may be more sensitive to parameter choices than others (Hleap et al., 2021). For example, we observed more overconfidence in ITS than COI, potentially reflecting a calibration more optimized for barcodes with lower base heterogeneity.

## Acknowledgements

We thank Duane D. McKenna, Renato J. P. Machado, Jia-Yong Zhang, and Shaun L. Winterton for generously sharing phylogenetic trees that served as backbone trees in our reference tree inference. The contributions of JO and TR were funded by the European Research Council (ERC) under the European Union’s Horizon 2020 research and innovation programme (grant agreement No 856506; ERC-synergy project LIFEPLAN). The computations were enabled by resources provided by the National Academic Infrastructure for Supercomputing in Sweden (NAISS), partially funded by the Swedish Research Council through grant agreement no. 2022-06725, and also by CSC - IT Center for Science, Finland.

## Author contributions

JO: Co-first author. Formal Analysis, Investigation, Visualization, Writing - original draft; AZ: Co-first author. Conceptualization, Methodology, Writing - Review & Editing; PS: Software, Writing - Review & Editing; JPH: Investigation, Writing - Review & Editing; MK: Investigation, Writing - Review & Editing; OO: Investigation, Writing - Review & Editing; MPB: Supervision, Writing - Review & Editing; NC: Investigation, Supervision, Writing - Review & Editing; TR: Supervision, Writing - Review & Editing; BF: Conceptualization, Investigation, Methodology, Software, Supervision, Writing - Review & Editing.

## Data availability

Data and code are available at https://github.com/jorsholm/taxclass.

