## Supplemental information for "Discovering the unseen: a performance comparison of taxonomic classification methods for unknown DNA barcodes"

### S1 Supplementary methods

#### S1.1 COI sequences

For COI sequences,only records identified to species level, not belonging to a "placeholder" taxon (here denoting the presence of "sp.", "aff.", "cf.", "nr.", "agg.", "t.", or "cluster" in the species name, or digits anywhere in the taxonomy string) were retained. Test and training sequences were aligned to the Folmer barcode region (Folmer et al., 1994) and amino acid translations were generated using MACSE v2.07 (Ranwez et al., 2018) and the COI coding sequence from *Drosophila melanogaster* (NCBI accession NC_024511.2, positions 1474..3009) as a reference. First, a set of 100 representative sequences from the training set were chosen using the MACSE "representative_seqs_v01" pipeline (Delsuc and Ranwez, 2020), using the COI coding sequence from *Drosophila melanogaster* (NCBI accession NC_024511.2, positions 1474..3009) as a reference. These representative sequences were then simultaneously aligned and translated with the MACSE `alignSequences` command using the invertebrate mitochondrial genetic code (option `-gc_def 5`) and terminal gap opening and extension penalties of 2.0 and 0.2, respectively (options `-gap_op_term 2.0` and "-gap_ext_term 0.2"), again using the *D. melanogaster* sequence as reference. The train and test datasets were then each aligned to the representative nucleotide and amino acid alignments and translated using the MACSE `enrichAlignment` command with the "-output_only_added_seq_ON" and "-fixed_alignment_ON" options, also removing any sequences with in-frame stop codons or internal insertions ("-maxSTOP_inSeq 0", "-maxINS_inSeq 0"), and using the same alignment penalties and genetic code as above. Columns containing more than 50% gaps, which included all positions outside the Folmer region which is most represented in the BOLD database, were removed using the esl-alimask command (distributed with HMMER3, Eddy, 2011) and trailing partial codons in the nucleotide alignments (one or two nucleotides separated from the rest of the sequence by one or two gaps) were rejoined to the main sequence using a custom awk script.

Alternate versions of the aligned and unaligned sequences with headers giving the taxonomic labels in several different header formats, as supported by the different algorithms, were generated using the Unix command line tools sed, tr, and awk in custom scripts.

#### S1.2 Classification algorithms

##### S1.2.1 Sequence similarity methods

We tested two separate approaches using BLAST as a taxonomic classifier: 1) top hit classification, where we accepted the taxonomy of the top hit as a prediction regardless of the sequence similarity; and 2) threshold classification, where we used predetermined similarity thresholds to accept or reject classifications at each taxonomic rank. To reflect common practice, we used for COI: class: 90%, order: 93%, family: 95%, subfamily: 96%, tribe: 96.5%, genus: 97%, and species: 98% (Ratnasingham, 2019), and for ITS: kingdom: 65%, phylum: 70%, class: 75%, order: 80%, family: 85%, genus: 90%, and species: 98% (Tedersoo et al., 2014). BLAST threshold generates missing predictions by discarding hits when sequence similarity is low, but BLAST top hit also generates missing predictions in some cases where there is no hit to the query sequence at all. We performed the thresholding operation during postprocessing the BLAST results, which used minimal computational resources, so separate listings for the top hit and threshold variants are not provided in Table S3.

We installed CREST4 version 4.3.7 from pip in a Python 3.9 virtual environment. Use of a custom reference database in CREST4 requires use of the separate script "make_new_crest_db.py" from crest4_utils, which we downloaded from `https://github.com/xapple/crest4_utils.git`. We discovered and corrected several minor bugs in crest4_utils during our tests. The patched version used for our tests is available at `githubreporemovedforanonymity`. We found that CREST4 did not assign the highest-level taxonomic rank provided in the reference database, so to obtain class-level assignments we added the phylum "Arthropoda" to the taxonomic annotation of all reference sequences for the COI train dataset before use with CREST4. No such change was required for the ITS dataset, which already included kingdom "Fungi" for all sequences. Hit lists were generated using BLAST 2.15.0 with recommended options from the CREST4 documentation (in particular `-num_alignments 100`), and then processed into classifications using the `crest4` command.

We downloaded dnabarcoder version 1.0.7 from `https://github.com/vuthuyduong/dnabarcoder`, and ran it in a Python 3.9 virtual environment initialized with biopython and matplotlib. Distance calculations were performed using `dnabarcoder.py sim`. Optimal thresholds were then performed using two calls per rank to `dnabarcoder.py predict`; first once at each rank to calculate global thresholds for that rank (e.g., `-rank family`), and then again to calculate local thresholds using the `-superrank` argument to specify all higher ranks (e.g., `-rank family -superrank order,class,phylum,kingdom`). For both global and local thresholds at all ranks we tested similarity thresholds from 0.85 to 1.0 in steps of 0.001 (`-st 0.85 -et 1.0 -s 0.001`). We then combined all the global and local thresholds into a single database using `dnabarcoder.py best`. We then generated hit lists for the test datasets using `dnabarcoder.py search`, and processed the hits into classifications using `dnabarcoder.py classify`.

CREST4 and dnabarcoder both generate missing predictions.

###### S1.2.2 K-mer composition methods

For RDP-NBC, we considered version 2.14 (2023), downloaded from `https://sourceforge.net/projects/rdp-classifier/files/rdp-classifier/` and run using OpenJDK version 21. We used the `train` command to generate the trained model parameters, and added a renamed version of `src/data/classifier/16srrna/rRNAClassifier.properties` from the downloaded source code. We then used the `classify` command with confidence threshold 0 (`-conf 0`) to output all classifications regardless of confidence limit for the FP setting.

For the MP setting, we removed classifications below the suggested confidence threshold of 0.8. RDP-NBC does not include any multithreading capability, so tests were executed only with a single CPU.

For SINTAX, we used the implementation in VSEARCH version 2.28.1 (Rognes et al., 2016), which made several minor changes to the algorithm to prevent incorrectly high confidence in the presence of ties. The model was initialized by creating an index using the `vsearch -makeudb-usearch` command, and predictions were made with `vsearch -sintax`, with the `-sintax-random` option. As for RDP-NBC, we removed classifications with confidence lower than 0.8 in the MP setting.

For IDTAXA, we installed DECIPHER 2.26.0 using renv 1.0.4 in R 4.2.3. Models were trained using the `LearnTaxa()` command with default parameters. Predictions were made using the `IdTaxa()` command with arguments `type=“collapsed”` for condensed output, `strand=“top”` to assume the input data are properly oriented, and `threshold=1` (meaning 1%) to output even low-confidence predictions. Because IDTAXA operates hierarchically, avoiding classification at lower ranks in low-confidence cases, it is likely that the low threshold resulted in somewhat slower performance than typical usage. For the MP setting, we removed classifications below the suggested confidence threshold of 0.6 for IDTAXA.

##### S1.2.3 Probabilistic models with novelty

We considered different implementations of PROTAX for COI and ITS barcodes. For COI, we used the model formulation and codebase for aligned sequences (ProtaxA), as in FinPROTAX (Roslin et al., 2022). Source code and scripts were downloaded from https://github.com/psomervuo/protaxA (commit dff9be3, dated 24 Aug, 2023). Input taxonomies were generated using a custom `GAWK` script, based on the training labels for the base case, or on the full Arthropod taxonomy from BOLD snapshot 29-Mar-2024 (Ratnasingham et al., 2024; BOLD Systems, 2024) for the augmented case. We weighted the “unknown” taxa so that, at each taxonomic rank, the total weight for “unknown” taxa was equal to 5% of the total weight (Abarenkov et al., 2018). Models were trained using the training scripts provided in the “readme.txt” file in ProtaxA, with modifications for the number of taxonomic levels. Classification was performed using the compiled “classify_v2” executable which implements fast Hamming distance calculations for aligned sequences but includes no option for multithreading. Thus ProtaxA tests were executed only with a single CPU. For ITS, we used the version available at https://github.com/psomervuo/PROTAX, which uses USEARCH (Edgar, 2010) to calculate pairwise distances. Models were trained using modified versions of the scripts in “step1.txt” and “step2.txt” scripts, and predictions were made using the script in “step4.txt”. Scripts for both versions of PROTAX were run using Perl 5.26.3 and R 4.2.3. Markov chain Monte Carlo traces were evaluated and considered sufficiently converged after the default sampling, so no additional sampling was performed.

For BayesANT, we used the implementation available at https://github.com/alessandrozito/BayesANT. For the aligned COI sequences, we trained the model using the default multinomial kernel (option `typeseq = “aligned”, type_location = “single”`) paired with the maximum likelihood estimation of the Pitman-Yor parameters controlling novelty at each taxonomic level. For ITS, we considered a 5-mer decomposition of the fungal sequences using the option `typeseq = ‘‘not_aligned’’`.

##### S1.2.4 Phylogenetic methods

Phylogenetic placement algorithms typically require that the query and reference sequences are globally aligned, making them unsuitable for use with barcode regions for which multiple sequence alignment is not feasible, such as ITS. Therefore, we only tested this class of models for COI. We classified the test data using EPA-ng with three different reference trees: one generated entirely de-novo from the reference database with no constraints, one built with full taxonomic constraints, and one built with constraints based on previously established phylogenies for which relationships between families are resolved. We refer to the first case as the *free tree*, the second case as the *taxonomically constrained tree*, and the third case as the *phylogenetically constrained tree*.

For the free tree, we added the reference COI sequence of the nematode *Caenorhabditis elegans* (NCBI accession number NC_001328.1, positions 7845–9422) to the reference alignment as an outgroup. This was necessary because this method did not generate trees which consistently separated Arachnida from Insecta. The reference alignment was then used to calculate a tree by maximum likelihood search in IQTREE version 2.4.0 (Minh et al., 2020) with the GTR+G4 (generalized time reversible, with 4-category approximation to gamma-distributed rates) substitution model for the nucleotide alignment, and the mART+G4 (Arthropod mitochondrial substitution matrix, with 4-category approximation to gamma-distributed rates) for the amino acid alignment, using mode `-fast` in both cases for fast inference.

For the taxonomically constrained tree, we constructed a multifurcating taxonomic tree from the annotations of the training sequences using the command `prepare taxonomy-tree` in Gappa version 0.8.5. This tree was then supplied as a constraint (`-g`) for tree search in IQTREE, which was otherwise run as above.

For the phylogenetically constrained tree, we retrieved phylogenies resolving family relationships within the 19 arthropod orders included in the training data, along with phylogenies spanning Chelicerata and Insecta to use as backbone trees; see Table S2. All trees were rooted using outgroups from the original publications. Order and family classification of all genera in each tree were retrieved from BOLD (Ratnasingham et al., 2024; BOLD Systems, 2024) when present, otherwise from NCBI taxonomy (Schoch et al., 2020). These trees were then grafted to form a single combined tree, starting by grafting the Insecta tree at the LCA of insects in the Chelicerata tree, and then continuing by grafting each order at the LCA of that order within the combined tree. Outgroup taxa were removed from each order-level tree before it was grafted into the combined tree. We then pruned the combined tree to only contain families present in FinBOL, limiting to a single branch per family. In cases where a family was not monophyletic in the reference tree (31 of 472 families), the branch for that family was placed as a child of the LCA of that family in the original tree, as shown is Fig. S1. To generate a constraint tree, the training sequences belonging to each family were added as a polytomy on the branch for that family. 88 families in the training set were not represented in the backbone tree, so sequences belonging to these families were excluded from the initial alignment. From the subset alignment, a first reference tree was inferred with IQTREE 2.1.4 using mode `-fast`, substitution model GTR+G4, and the backbone tree to constrain the topology. To include the 88 families not represented in the initial backbone tree, the sequences belonging to these families were phylogenetically placed in the initial reference tree, using EPA-ng version 0.3.8. To find the most likely placement for each family, we calculated the product of likelihoods across all branches and placed sequences belonging to that family, identifying the maximum value to determine the optimal position. For three families (*Baetidae*, *Boreidae*, and *Diprionidae*), the most likely placement was within a different order than the family taxonomically belongs to, or forming a sister clade to a different order. These families were placed as children to the LCA of the correct order, that is, as a sister clade to all other sequences in that order. To create the final constraint tree, the 88 families were added at the position of their most likely placement in the first reference tree, and each family, including those in the initial reference tree, was again collapsed to a polytomy. The combined backbone tree was then supplied as a constraint for tree search in IQTREE, which was otherwise run as above.

In all three cases, we placed all query sequences into the reference phylogeny using EPA-ng version 0.3.8 and assigned taxonomic labels with Gappa version 0.8.5 (`examine assign`). The free tree was rooted in Gappa using the added *C. elegans* reference sequence, while the taxonomically and phylogenetically constrained trees were rooted using Arachnida.

###### S1.2.5 Neural network methods

There have been a number of studies applying neural-network models to taxonomic identification of DNA barcodes (e.g., Vu et al., 2020; Flück et al., 2022; Yang et al., 2022; Arias et al., 2023). However, most of these did not meet our requirements of providing classifications at each level of the taxonomic hierarchy, and including code and instructions to train and apply the model with user-provided reference data. We were thus able to test only two models in this category, both implemented in the same software package.

We installed MycoAI v0.0.5 from pypi (package name mycoai-its). We installed the package in a Python 3.9.7 virtual environment using pip 20.2.4. We used PyTorch 1.12, the minimum version compatible with MycoAI. Models were trained using `mycoai-train` from the command line. For the CNN model, the option `-base_arch_type CNN` was given. Test data were classified using `mycoai-classify` with option `-confidence` to output confidence scores.

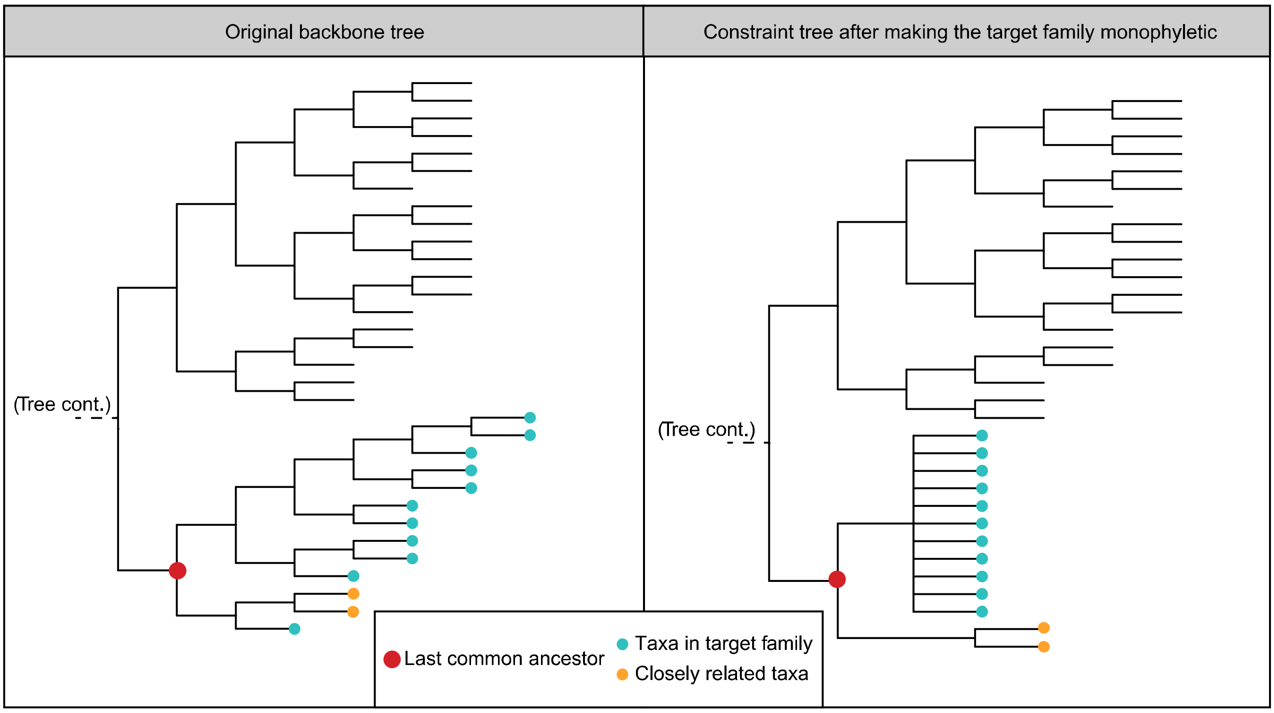


Figure S1: Illustration of how the constraint tree was edited to ensure monophyletic families. In cases where a family was not monophyletic in the original tree, the branch for that family was placed as child of the last common ancestor of that family in the original tree.

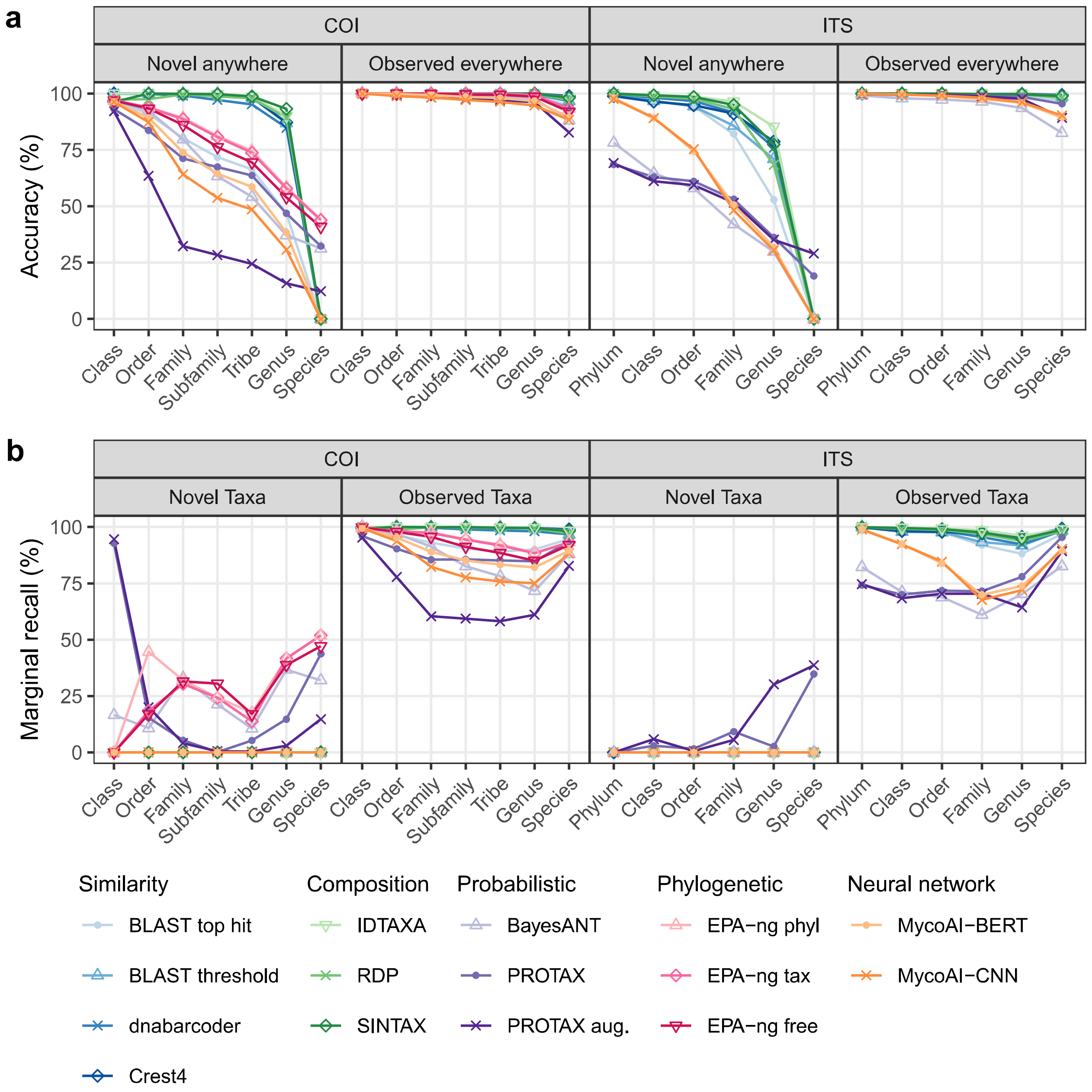


Figure S2: Accuracy and marginal recall of classification algorithms across taxonomic ranks under the MP setting, i.e. where predictions were enforced for all query sequences. **a** shows accuracy, i.e. the proportion of all taxonomic classifications that were correct for observed (right) and novel (left) species. **b** shows marginal recall, i.e. the proportion of correct predictions relative to the total number of sequences belonging to that respective taxon set and rank for observed (right) and novel (left) taxa.

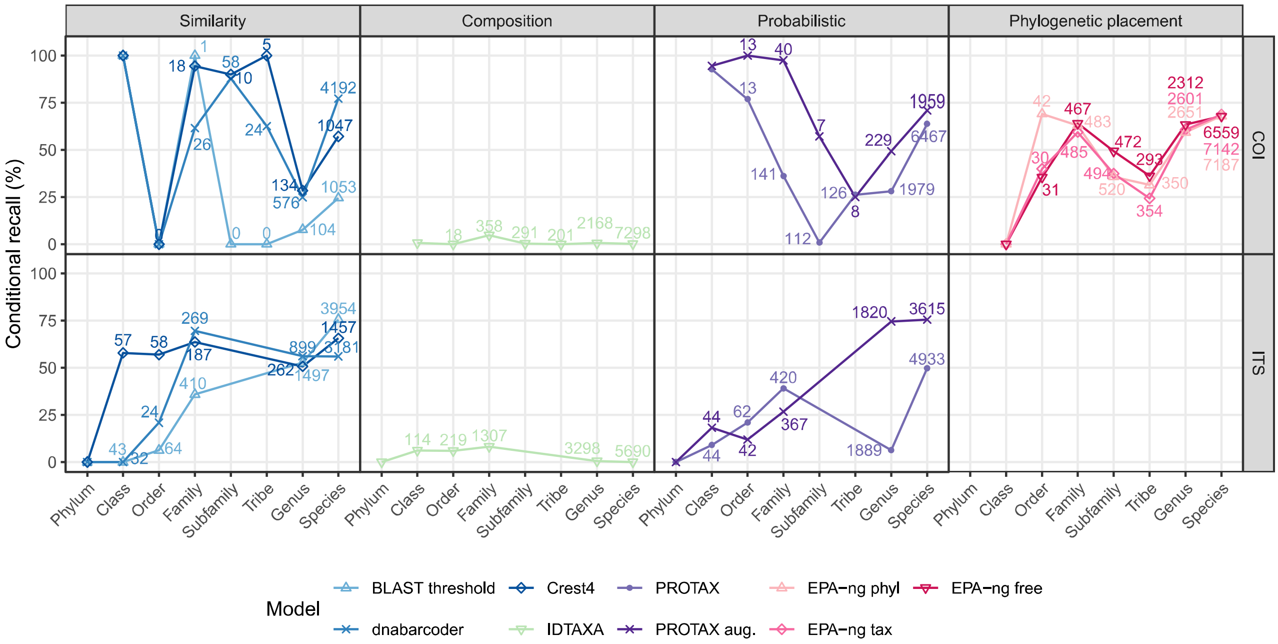


Figure S3: Conditional recall for taxonomic predictions of novel taxa. For each rank, it shows the proportion of correct predictions given that the rank above was classified correctly. Each algorithm is represented by a different shape and color, with hue indicating algorithm group: sequence similarity (blue), composition (green), probabilistic (purple), phylogenetic (pink), and neural network (orange). Numbers display the count of correct predictions on the rank above for each algorithm and rank, i.e., the number of predictions included in the denominator. For the highest rank, the denominator was 439 and 146 for COI and ITS, respecitvely.

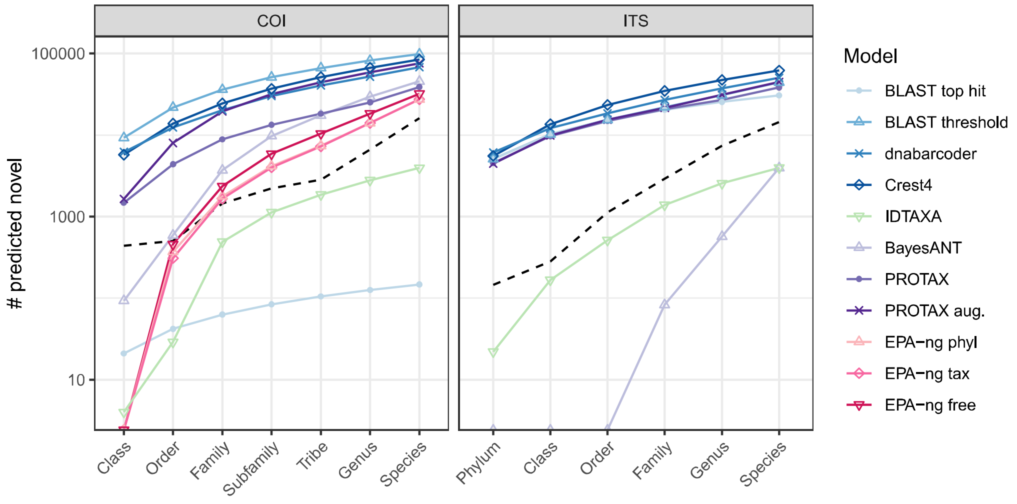


Figure S4: Cumulative number of taxa predicted as novel across ranks and classification algorithms. The true number of novel taxa is shown with a black dashed line. Only algorithms which explicitly predict novel taxa or produce missing predictions are included in the plot.

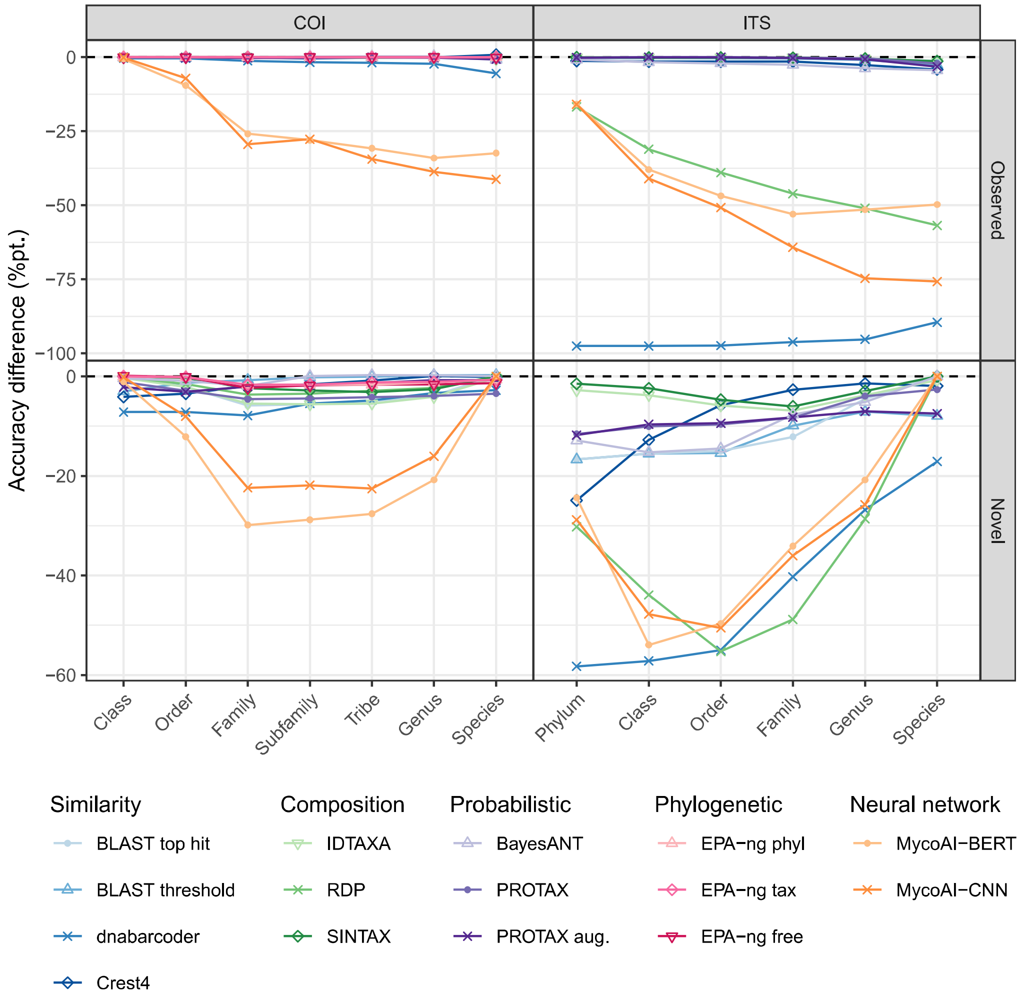


Figure S5: Change in classification accuracy when using full-length vs. "testshort" DNA sequences, shown in percentage points. Line of zero difference shown with a black dashed line. Negative values indicate lower accuracy of short reads than long reads, and vice versa.

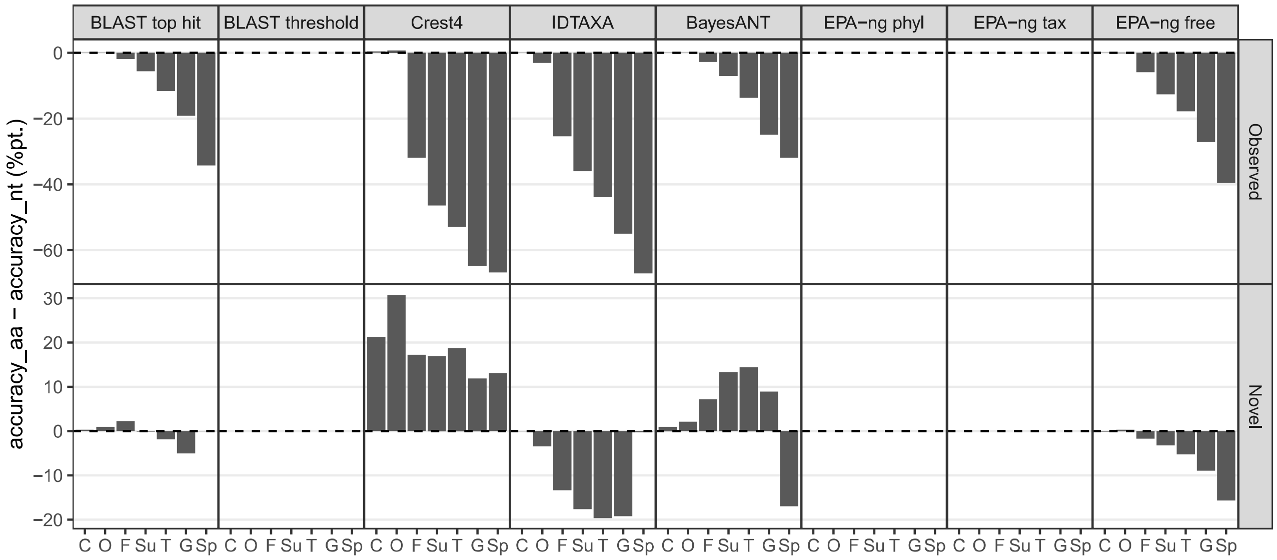


Figure S6: Change in classification accuracy when using nucleotide sequences vs. amino acid translation of the COI barcode, shown in percentage points. Line of zero difference shown with a black dashed line. Negative values indicate lower accuracy of amino acid sequences, and vice versa.

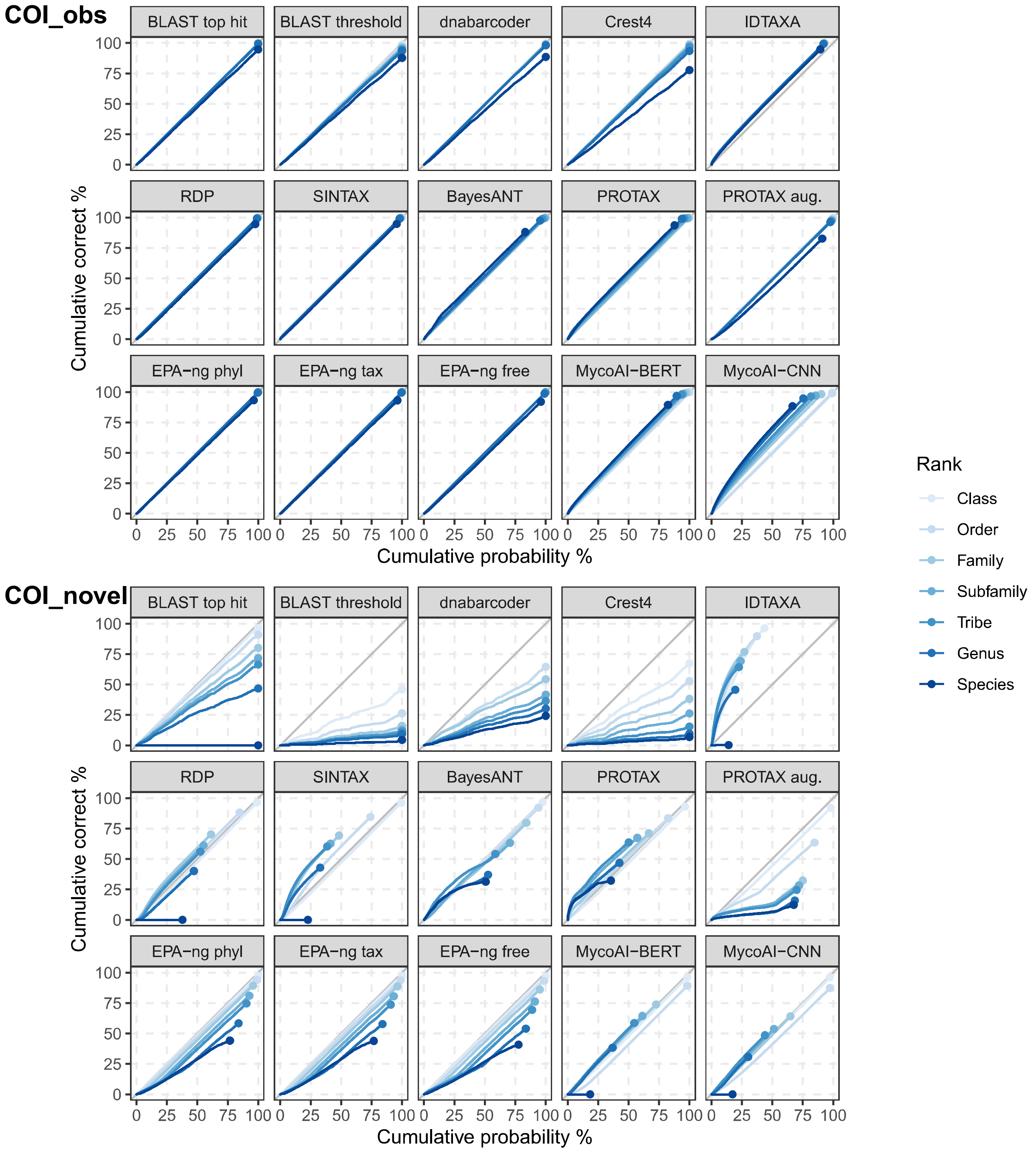

COI_obs
COI_novel
BLAST top hit
BLAST threshold
dnabarcoder
Crest4
IDTAXA
RDP
SINTAX
BayesANT
PROTAX
PROTAX aug.
EPA−ng phyl
EPA−ng tax
EPA−ng free
MycoAI−BERT
MycoAI−CNN
Cumulative correct %
Cumulative probability %
0
25
50
75
100
Rank
Class
Order
Family
Subfamily
Tribe
Genus
Species


Figure S7

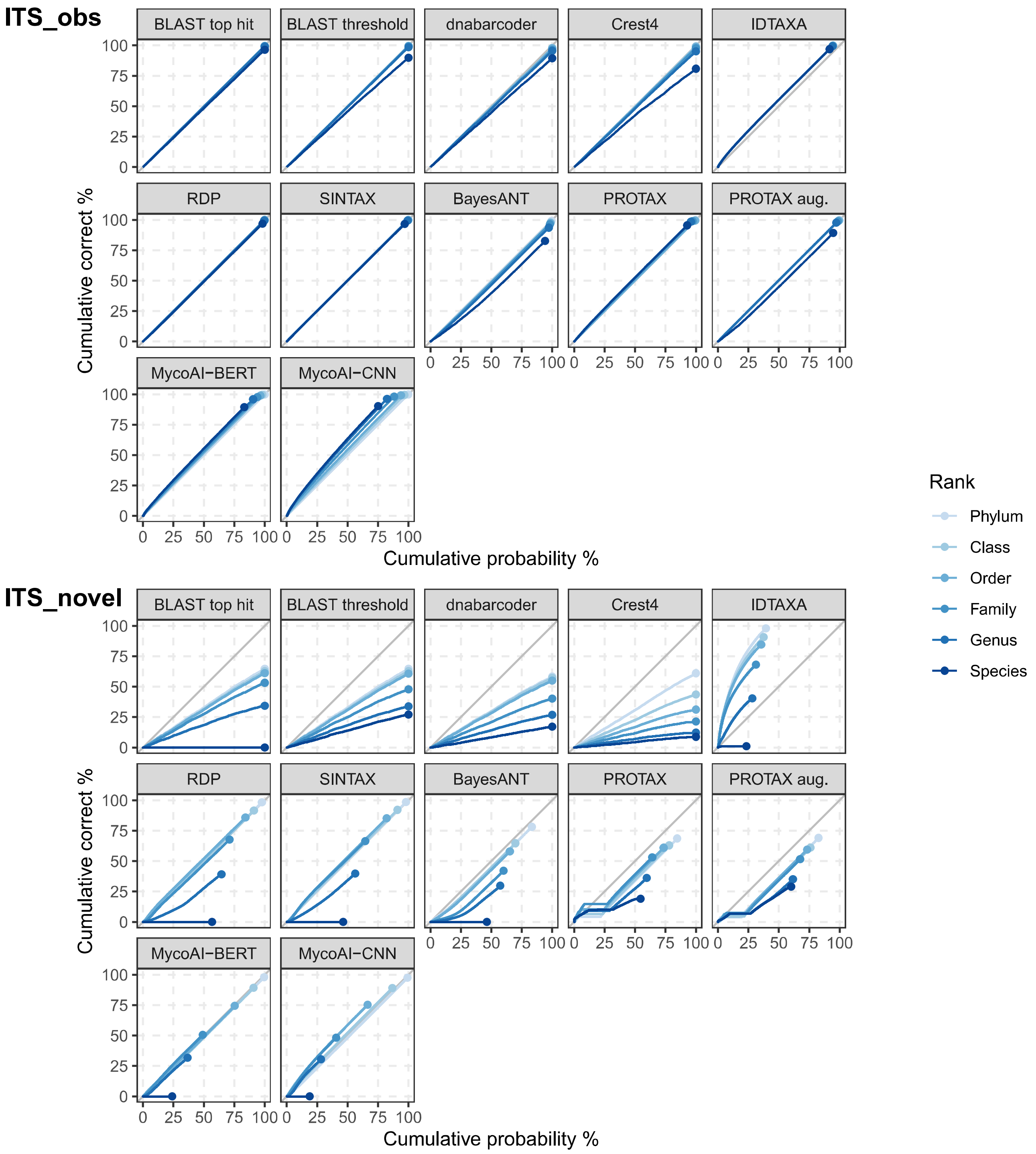

ITS_obs
BLAST top hit
BLAST threshold
dnabarcoder
Crest4
IDTAXA
RDP
SINTAX
BayesANT
PROTAX
PROTAX aug.
MycoAI−BERT
MycoAI−CNN
Cumulative correct %
Cumulative probability %
ITS_novel
BLAST top hit
BLAST threshold
dnabarcoder
Crest4
IDTAXA
RDP
SINTAX
BayesANT
PROTAX
PROTAX aug.
MycoAI−BERT
MycoAI−CNN
Cumulative correct %
Cumulative probability %
0
25
50
75
100
Rank
Phylum
Class
Order
Family
Genus
Species


Figure S8

### 5 Computational resources

Table S3: Computational resource use of different classification algorithms. Initialization/training and classification phases are shown separately for COI and ITS sequences. Reported metrics include wall-clock time, average CPU usage, peak memory consumption, and model size on disk. Values in parentheses are for classification of short sequences representative of metabarcoding data. Results are provided for both single-core and multi-core CPU runs, as well as GPU runs where available.

| Marker | Algorithm | CPU / GPU | Initialization/Training Time | CPU (%) | Mem (kB) | Classification Time | CPU (%) | Mem (kB) | Disk (kB) |
|---|---|---|---|---|---|---|---|---|---|
| COI NT | BayesANT | 1 | 4:40 | 89% | 1 539 720 | 1:35:31<br>(1:03:53) | 98%<br>(99%) | 933 968<br>(904 316) | 155 244 |
| COI NT | BayesANT | 4 | 3:07 | 92% | 1 537 884 | 18:52<br>(16:24) | 396%<br>(395%) | 917 460<br>(884 248) | 155 244 |
| COI NT | BayesANT | 16 | 3:27 | 91% | 1 533 648 | 6:06<br>(4:08) | 1472%<br>(1458%) | 917 452<br>(884 624) | 155 244 |
| COI NT | BayesANT | 40 | 3:04 | 90% | 1 537 696 | 2:43<br>(1:53) | 3763%<br>(3686%) | 917 384<br>(883 704) | 155 244 |
| COI AA | BayesANT | 1 | 2:37 | 95% | 2 238 540 | 41:18<br>(28:17) | 99%<br>(99%) | 1 261 320<br>(1 248 048) | 237 896 |
| COI AA | BayesANT | 4 | 2:35 | 94% | 2 236 132 | 10:50<br>(7:38) | 388%<br>(385%) | 1 248 092<br>(1 236 952) | 237 896 |
| COI AA | BayesANT | 16 | 1:49 | 98% | 2 236 932 | 2:43<br>(1:59) | 1469%<br>(1435%) | 1 248 212<br>(1 237 068) | 237 896 |
| COI AA | BayesANT | 40 | 1:45 | 98% | 2 236 952 | 1:12<br>(0:54) | 3542%<br>(3384%) | 1 247 924<br>(1 236 760) | 237 896 |
| COI NT | BLAST | 1 | 0:02 | 39% | 12 200 | 2:16:49<br>(48:20) | 98%<br>(98%) | 44 944<br>(49 080) | 13 212 |
| COI NT | BLAST | 4 | 0:02 | 33% | 12 860 | 43:47<br>(20:23) | 391%<br>(320%) | 148 600<br>(122 776) | 13 212 |
| COI NT | BLAST | 16 | 0:02 | 33% | 12 316 | 31:25<br>(17:05) | 445%<br>(321%) | 146 580<br>(136 968) | 13 212 |
| COI NT | BLAST | 40 | 0:02 | 29% | 12 620 | 29:27<br>(15:60) | 446%<br>(319%) | 155 680<br>(127 116) | 13 212 |
| COI AA | BLAST | 1 | 0:02 | 37% | 12 224 | 10:50:39<br>(8:10:41) | 99%<br>(97%) | 81 692<br>(115 572) | 14 952 |
| COI AA | BLAST | 4 | 0:02 | 35% | 12 204 | 2:40:34<br>(1:51:53) | 393%<br>(394%) | 244 612<br>(368 796) | 14 952 |
| COI AA | BLAST | 16 | 0:01 | 39% | 12 200 | 44:11<br>(33:20) | 1566%<br>(1559%) | 894 152<br>(1 384 580) | 14 952 |
| COI AA | BLAST | 40 | 0:01 | 38% | 12 172 | 17:54<br>(13:55) | 3854%<br>(3710%) | 2 193 584<br>(3 429 804) | 14 952 |

| | | | | | | | | | |
|---|---|---|---|---|---|---|---|---|---|
| COI NT | CREST4 | 1 | 0:12 | 18% | 56 788 | 2:35:27 (1:07:11) | 98% (98%) | 4 127 012 (4 103 068) | 47 956 |
| COI NT | CREST4 | 4 | 0:13 | 16% | 58 672 | 1:00:13 (35:18) | 337% (256%) | 4 128 272 (4 102 996) | 47 952 |
| COI NT | CREST4 | 16 | 0:12 | 28% | 56 956 | 42:36 (32:04) | 377% (254%) | 4 130 872 (4 104 580) | 47 952 |
| COI NT | CREST4 | 40 | 0:11 | 60% | 58 624 | 38:55 (25:45) | 377% (255%) | 4 131 872 (4 112 980) | 47 952 |
| COI AA | CREST4 | 1 | 0:04 | 58% | 56 924 | 12:07:41 (9:05:44) | 99% (99%) | 3 968 084 (3 964 700) | 40 040 |
| COI AA | CREST4 | 4 | 0:03 | 70% | 57 056 | 3:26:17 (2:34:16) | 379% (377%) | 3 969 408 (3 964 704) | 40 040 |
| COI AA | CREST4 | 16 | 0:03 | 141% | 60 352 | 55:35 (42:56) | 1398% (1302%) | 3 977 328 (3 964 904) | 40 044 |
| COI AA | CREST4 | 40 | 0:04 | 148% | 61 444 | 25:56 (21:34) | 2987% (2681%) | 3 975 060 (3 972 556) | 40 040 |
| COI NT | dnabarcoder | 1 | 6:49:29 | 93% | 2 477 792 | 3:08:03 (1:27:39) | 99% (99%) | 1 542 756 (1 682 908) | 564 |
| COI NT | dnabarcoder | 4 | 2:18:55 | 235% | 50 536 572 | 46:37 (26:02) | 383% (320%) | 1 476 036 (1 500 260) | 632 |
| COI NT | dnabarcoder | 16 | 2:20:04 | 246% | 50 533 764 | 41:05 (27:07) | 447% (321%) | 1 456 528 (1 551 204) | 632 |
| COI NT | dnabarcoder | 40 | 1:58:41 | 257% | 50 542 992 | 38:19 (25:12) | 441% (319%) | 1 516 304 (1 690 124) | 632 |
| COI NT | EPA-ng free | 1 | 4:03:24 | 99% | 38 356 188 | 34:14 (28:12) | 99% (99%) | 36 714 264 (30 745 376) | 14 524 |
| COI NT | EPA-ng free | 4 | 4:49:57 | 292% | 38 353 036 | 14:23 (11:53) | 357% (351%) | 36 597 496 (30 731 504) | 14 644 |
| COI NT | EPA-ng free | 16 | 4:50:44 | 932% | 38 382 320 | 4:34 (3:53) | 1085% (1054%) | 36 654 012 (30 799 696) | 14 356 |
| COI NT | EPA-ng free | 40 | 2:52:44 | 3454% | 38 388 548 | 2:29 (2:10) | 2126% (1978%) | 36 672 944 (30 779 940) | 15 668 |
| COI AA | EPA-ng free | 1 | 2d 04:53:01 | 99% | 41 795 608 | 1d 22:06:19 (1d 12:04:35) | 99% (99%) | 39 414 660 (32 864 812) | 17 600 |
| COI AA | EPA-ng free | 4 | 14:38:14 | 375% | 41 794 392 | 12:57:54 (14:04:55) | 397% (397%) | 39 501 880 (33 167 828) | 20 172 |
| COI AA | EPA-ng free | 16 | 5:42:06 | 1339% | 41 794 908 | 3:06:57 (3:10:43) | 1576% (1578%) | 39 644 404 (33 152 212) | 17 752 |
| COI AA | EPA-ng free | 40 | 4:03:38 | 3096% | 41 800 640 | 1:21:13 (1:23:40) | 3888% (3901%) | 39 660 140 (33 162 612) | 21 488 |
| COI NT | EPA-ng phyl | 1 | 13:32:55 | 96% | 4 046 284 | 50:08 (39:58) | 98% (98%) | 36 586 616 (30 701 520) | 14 900 |
| COI NT | EPA-ng phyl | 4 | 11:38:43 | 172% | 4 732 332 | 10:11 | 360% | 36 631 416 | 15 428 |

| | | | | | | | | | |
|---|---|---|---|---|---|---|---|---|---|
| | | | | | | (8:17) | (352%) | (30 744 412) | |
| COI NT | EPA-ng phyl | 16 | 19:13:36 | 369% | 7 477 204 | 5:27<br>(4:06) | 916%<br>(970%) | 36 649 392<br>(30 790 020) | 15 220 |
| COI NT | EPA-ng phyl | 40 | 10:51:58 | 1274% | 12 965 244 | 2:31<br>(2:09) | 2001%<br>(1853%) | 36 657 072<br>(30 784 000) | 15 216 |
| COI AA | EPA-ng phyl | 1 | 10:27:50 | 99% | 7 580 728 | 2d 02:11:06<br>(2d 01:00:36) | 99%<br>(99%) | 39 408 124<br>(32 781 344) | 15 780 |
| COI AA | EPA-ng phyl | 4 | 10:39:31 | 146% | 8 267 936 | 11:57:10<br>(13:10:52) | 397%<br>(397%) | 39 510 104<br>(33 135 284) | 15 436 |
| COI AA | EPA-ng phyl | 16 | 14:02:13 | 285% | 10 670 080 | 3:16:29<br>(3:23:16) | 1565%<br>(1571%) | 39 576 340<br>(33 134 068) | 16 360 |
| COI AA | EPA-ng phyl | 40 | 10:32:23 | 519% | 16 500 224 | 1:30:04<br>(1:22:51) | 3897%<br>(3885%) | 39 671 800<br>(33 182 440) | 16 628 |
| COI NT | EPA-ng taxa | 1 | 12:31:42 | 99% | 4 039 520 | 50:00<br>(36:38) | 99%<br>(95%) | 36 588 184<br>(30 703 300) | 14 884 |
| COI NT | EPA-ng taxa | 4 | 11:51:53 | 182% | 4 717 756 | 12:47<br>(10:31) | 365%<br>(361%) | 36 594 500<br>(30 696 512) | 12 756 |
| COI NT | EPA-ng taxa | 16 | 18:53:08 | 386% | 7 434 196 | 4:48<br>(4:01) | 1043%<br>(1018%) | 36 633 564<br>(30 791 788) | 14 132 |
| COI NT | EPA-ng taxa | 40 | 10:37:17 | 1322% | 12 873 192 | 2:29<br>(2:10) | 2107%<br>(1934%) | 36 663 996<br>(30 749 088) | 12 740 |
| COI AA | EPA-ng taxa | 1 | 10:51:36 | 99% | 7 574 524 | 1d 23:58:25<br>(1d 21:51:03) | 99%<br>(99%) | 39 441 232<br>(32 885 796) | 18 768 |
| COI AA | EPA-ng taxa | 4 | 12:06:34 | 146% | 8 256 200 | 12:58:55<br>(13:42:18) | 397%<br>(397%) | 39 633 692<br>(33 087 680) | 16 412 |
| COI AA | EPA-ng taxa | 16 | 15:01:38 | 319% | 10 970 504 | 3:17:10<br>(3:21:50) | 1573%<br>(1574%) | 39 578 588<br>(33 147 028) | 14 344 |
| COI AA | EPA-ng taxa | 40 | 11:15:06 | 558% | 16 402 824 | 1:22:58<br>(1:24:10) | 3887%<br>(3898%) | 39 680 484<br>(33 201 092) | 14 092 |
| COI NT | IDTAXA | 1 | 14:46 | 97% | 2 535 520 | 2:13:46<br>(1:49:21) | 99%<br>(99%) | 1 001 704<br>(992 264) | 50 684 |
| COI NT | IDTAXA | 4 | 14:53 | 97% | 2 570 152 | 42:13<br>(34:36) | 342%<br>(335%) | 1 028 316<br>(992 756) | 50 680 |
| COI NT | IDTAXA | 16 | 12:05 | 98% | 2 478 764 | 16:15<br>(13:40) | 992%<br>(964%) | 1 010 700<br>(1 001 428) | 50 680 |
| COI NT | IDTAXA | 40 | 11:12 | 97% | 2 556 068 | 12:06<br>(10:22) | 1685%<br>(1630%) | 1 010 036<br>(1 023 592) | 50 680 |
| COI AA | IDTAXA | 1 | 53:10 | 97% | 12 506 364 | 43:51<br>(22:49) | 99%<br>(99%) | 842 960<br>(751 760) | 4 788 |
| COI AA | IDTAXA | 4 | 54:16 | 99% | 12 808 980 | 15:42<br>(9:05) | 305%<br>(286%) | 813 140<br>(839 864) | 4 788 |
| COI AA | IDTAXA | 16 | 43:59 | 99% | 12 962 520 | 8:10<br>(4:42) | 731%<br>(748%) | 821 412<br>(795 744) | 4 788 |

| | | | | | | | | | |
|---|---|---|---|---|---|---|---|---|---|
| COI AA | IDTAXA | 40 | 40:40 | 99% | 12 477 072 | 6:14 (4:06) | 1395% (1367%) | 813 796 (804 676) | 4 788 |
| COI NT | MycoAI-BERT | 1 | 5d 13:49:00 | 96% | 11 079 492 | 26:44 (14:39) | 97% (99%) | 2 167 976 (3 649 280) | 438 600 |
| COI NT | MycoAI-BERT | 4 | 1d 18:09:02 | 396% | 11 294 900 | 15:06 (6:32) | 280% (389%) | 2 872 980 (21 525 648) | 424 740 |
| COI NT | MycoAI-BERT | 16 | 14:25:40 | 1568% | 11 464 400 | 5:03 (2:34) | 1233% (1387%) | 2 551 460 (3 792 064) | 418 672 |
| COI NT | MycoAI-BERT | 40 | 10:18:38 | 3840% | 11 925 688 | 3:02 (1:27) | 3127% (3608%) | 2 289 256 (4 006 960) | 422 164 |
| COI NT | MycoAI-BERT | GPU | 43:41 | 95% | 3 183 728 | 1:07 (0:31) | 60% (95%) | 2 433 940 (2 421 804) | 439 148 |
| COI NT | MycoAI-CNN | 1 | 4:00:39 | 98% | 1 115 268 | 1:38 (0:32) | 29% (73%) | 1 498 156 (794 808) | 202 284 |
| COI NT | MycoAI-CNN | 4 | 1:28:20 | 370% | 1 187 432 | 1:01 (0:24) | 64% (142%) | 1 194 224 (1 716 256) | 202 856 |
| COI NT | MycoAI-CNN | 16 | 37:58 | 1364% | 1 284 800 | 0:54 (0:23) | 93% (189%) | 1 839 624 (842 872) | 203 396 |
| COI NT | MycoAI-CNN | 40 | 34:03 | 3154% | 1 577 352 | 0:20 (0:18) | 475% (500%) | 1 775 912 (1 783 064) | 203 240 |
| COI NT | MycoAI-CNN | GPU | 9:19 | 85% | 3 787 128 | 0:27 (0:23) | 95% (94%) | 3 175 540 (3 184 600) | 202 460 |
| COI NT | ProtaxA | 1 | 4:35:12 | 98% | 426 924 | 1:28 (1:27) | 99% (99%) | 104 320 (104 288) | 373 200 |
| COI NT | RDP-NBC | 1 | 0:30 | 88% | 3 859 068 | 18:49 (10:13) | 99% (99%) | 1 122 064 (1 125 116) | 122 828 |
| COI NT | SINTAX | 1 | 0:09 | 86% | 79 676 | 7:11 (7:01) | 99% (99%) | 123 248 (123 160) | 112 360 |
| COI NT | SINTAX | 4 | 0:08 | 94% | 80 888 | 1:53 (1:50) | 384% (390%) | 123 484 (125 372) | 112 360 |
| COI NT | SINTAX | 16 | 0:07 | 84% | 80 076 | 1:14 (1:19) | 1468% (1457%) | 141 768 (145 704) | 112 360 |
| COI NT | SINTAX | 40 | 0:06 | 91% | 79 856 | 1:14 (1:14) | 3850% (3850%) | 142 608 (146 896) | 112 360 |
| ITS NT | BayesANT | 1 | 0:36 | 68% | 884 044 | 2:00:18 (2:03:09) | 99% (99%) | 1 181 204 (1 092 664) | 30 064 |
| ITS NT | BayesANT | 4 | 0:37 | 63% | 885 392 | 28:04 (28:01) | 395% (395%) | 1 181 484 (1 088 116) | 30 064 |
| ITS NT | BayesANT | 16 | 0:56 | 33% | 887 288 | 8:41 (8:57) | 1538% (1552%) | 1 372 324 (1 087 620) | 30 068 |
| ITS NT | BayesANT | 40 | 0:36 | 51% | 894 472 | 3:52 (3:46) | 3691% (3745%) | 1 372 000 (1 084 720) | 30 064 |

| | | | | | | | | | |
|---|---|---|---|---|---|---|---|---|---|
| ITS NT | BLAST | 1 | 0:02 | 18% | 12 164 | 30:18<br>(0:26) | 99%<br>(97%) | 39 144<br>(1 492 316) | 3 688 |
| ITS NT | BLAST | 4 | 0:01 | 15% | 12 096 | 6:35<br>(0:08) | 386%<br>(260%) | 41 924<br>(1 554 928) | 3 684 |
| ITS NT | BLAST | 16 | 0:02 | 12% | 12 084 | 2:06<br>(0:05) | 1482%<br>(468%) | 46 720<br>(1 654 348) | 3 688 |
| ITS NT | BLAST | 40 | 0:02 | 11% | 12 132 | 1:45<br>(0:05) | 1807%<br>(471%) | 53 860<br>(1 773 384) | 3 688 |
| ITS NT | CREST4 | 1 | 0:12 | 7% | 41 936 | 39:21<br>(2:24) | 99%<br>(99%) | 2 611 660<br>(2 155 132) | 12 176 |
| ITS NT | CREST4 | 4 | 0:15 | 8% | 41 860 | 14:20<br>(1:59) | 273%<br>(119%) | 2 611 780<br>(2 266 384) | 12 176 |
| ITS NT | CREST4 | 16 | 0:11 | 24% | 40 956 | 6:55<br>(1:31) | 564%<br>(131%) | 2 612 312<br>(2 494 464) | 12 176 |
| ITS NT | CREST4 | 40 | 0:13 | 41% | 41 288 | 6:54<br>(1:26) | 559%<br>(135%) | 2 612 200<br>(2 913 932) | 12 176 |
| ITS NT | dnabarcoder | 1 | 1d 01:29:45 | 98% | 23 450 900 | 47:21<br>(1:14) | 99%<br>(97%) | 360 412<br>(3 661 468) | 132 |
| ITS NT | dnabarcoder | 4 | 19:36:49 | 101% | 4 508 436 | 20:22<br>(0:54) | 256%<br>(158%) | 443 440<br>(3 671 756) | 132 |
| ITS NT | dnabarcoder | 16 | 16:20:51 | 101% | 4 504 840 | 8:37<br>(0:39) | 565%<br>(204%) | 329 544<br>(3 680 444) | 132 |
| ITS NT | dnabarcoder | 40 | 15:06:21 | 102% | 4 506 532 | 8:37<br>(0:33) | 554%<br>(243%) | 322 716<br>(3 688 992) | 132 |
| ITS NT | IDTAXA | 1 | 7:38 | 96% | 1 594 708 | 30:57<br>(13:06) | 99%<br>(99%) | 780 868<br>(707 608) | 12 296 |
| ITS NT | IDTAXA | 4 | 9:18 | 97% | 1 543 460 | 13:26<br>(8:17) | 291%<br>(259%) | 796 268<br>(709 336) | 12 296 |
| ITS NT | IDTAXA | 16 | 5:48 | 95% | 1 543 308 | 6:12<br>(4:00) | 831%<br>(844%) | 770 132<br>(708 648) | 12 296 |
| ITS NT | IDTAXA | 40 | 6:00 | 88% | 1 543 320 | 5:31<br>(3:47) | 1591%<br>(1839%) | 772 908<br>(710 788) | 12 296 |
| ITS NT | MycoAI-BERT | 1 | 2d 03:07:01 | 98% | 10 239 096 | 57:18<br>(55:08) | 95%<br>(98%) | 2 247 300<br>(2 213 640) | 308 012 |
| ITS NT | MycoAI-BERT | 4 | 17:20:58 | 381% | 10 177 120 | 15:47<br>(14:18) | 338%<br>(367%) | 2 255 304<br>(2 222 320) | 378 332 |
| ITS NT | MycoAI-BERT | 16 | 5:56:57 | 1489% | 10 493 340 | 7:60<br>(6:05) | 1126%<br>(1478%) | 2 275 948<br>(2 246 384) | 344 348 |
| ITS NT | MycoAI-BERT | 40 | 5:20:46 | 3596% | 10 801 436 | 7:48<br>(5:57) | 2710%<br>(3585%) | 2 289 216<br>(2 263 748) | 372 312 |
| ITS NT | MycoAI-BERT | GPU | 16:55 | 70% | 1 406 356 | 0:30<br>(0:27) | 93%<br>(95%) | 927 632<br>(918 776) | 292 680 |
| ITS NT | MycoAI-CNN | 1 | 24:52 | 81% | 687 324 | 0:19 | 92% | 605 376 | 37 236 |

| | | | | | | | | | |
|---|---|---|---|---|---|---|---|---|---|
| | | | | | | (0:16) | (91%) | (600 556) | |
| ITS NT | MycoAI-CNN | 4 | 10:10 | 236% | 696 832 | 0:15<br>(0:12) | 121%<br>(129%) | 607 192<br>(602 568) | 37 536 |
| ITS NT | MycoAI-CNN | 16 | 7:40 | 693% | 698 404 | 0:13<br>(0:10) | 217%<br>(245%) | 605 804<br>(601 008) | 37 308 |
| ITS NT | MycoAI-CNN | 40 | 8:51 | 1429% | 708 744 | 0:13<br>(0:10) | 444%<br>(547%) | 606 360<br>(601 576) | 37 200 |
| ITS NT | MycoAI-CNN | GPU | 7:43 | 47% | 1 354 944 | 0:22<br>(0:17) | 86%<br>(92%) | 816 624<br>(812 044) | 37 552 |
| ITS NT | Protax | 1 | 3:12:18 | 99% | 373 084 | 8:34<br>(4:26) | 99%<br>(99%) | 198 292<br>(123 644) | 199 932 |
| ITS NT | Protax | 4 | 4:04:12 | 100% | 373 028 | 5:09<br>(3:23) | 219%<br>(175%) | 198 332<br>(123 684) | 199 916 |
| ITS NT | Protax | 16 | 3:16:05 | 101% | 372 984 | 3:18<br>(2:27) | 311%<br>(219%) | 198 292<br>(123 624) | 199 928 |
| ITS NT | Protax | 40 | 2:27:59 | 101% | 372 836 | 2:45<br>(2:01) | 299%<br>(211%) | 198 264<br>(123 624) | 199 896 |
| ITS NT | RDP-NBC | 1 | 0:16 | 68% | 1 614 400 | 3:16<br>(0:51) | 97%<br>(99%) | 720 780<br>(730 780) | 35 876 |
| ITS NT | SINTAX | 1 | 0:02 | 77% | 26 644 | 1:09<br>(0:56) | 99%<br>(99%) | 33 036<br>(32 996) | 27 776 |
| ITS NT | SINTAX | 4 | 0:01 | 99% | 26 840 | 0:22<br>(0:19) | 378%<br>(367%) | 33 360<br>(33 220) | 27 776 |
| ITS NT | SINTAX | 16 | 0:01 | 97% | 26 760 | 0:42<br>(0:43) | 1484%<br>(1481%) | 34 028<br>(34 072) | 27 776 |
| ITS NT | SINTAX | 40 | 0:02 | 69% | 26 672 | 0:48<br>(0:48) | 3865%<br>(3860%) | 35 824<br>(35 788) | 27 776 |

Table S1: Additional algorithms for taxonomic classification and why they were not included in our evaluation.

| Algorithm | Citation | Short description | Reason for exclusion |
|---|---|---|---|
| RTAX | Soergel et al. (2012) | Developed for classification of pair-end sequences. | Poor fit for the concatenated sequences in this comparison. |
| UTAX | Edgar (Edgar) | Original algorithm in USEARCH before SINTAX. | Author suggest using SINTAX instead. |
| Mycofier | Delgado-Serrano et al. (2016) | Naïve Bayesian classifier based on 5-mers, sequence length, and GC content. | Default database gives only species output. |
| MEGAN | Huson et al. (2016) | LCA classifier. | Constrained to NCBI taxonomy. |
| Qiime2 | Bokulich et al. (2018) | Creates consensus taxonomy from RDP, SINTAX, and BLAST classifications. | |
| CONSTAX | Gdanetz et al. (2017); Liber et al. (2021) | | |
| funbarRF | Meher et al. (2019) | Random forest algorithm using base pair composition allowing gaps. | R package removed fron CRAN repository. |
| CNN | Helaly et al. (2020) | Different convolutional neural network (CNN) architectures and feature representations. | No code available with publication. |
| CNN & DBN | Vu et al. (2020) | CNN and deep belief network (DBN) developed for fungi. | Models predict only a single taxonomic rank. |
| Its2Vec | Wang et al. (2020) | Random-forest based classifier developed for ITS barcodes. | RAR file containing code could no be opened. |
| CNN | Flück et al. (2022) | CNN-based classifier developed for short fish barcodes (60 bp). | Unclear how it would work with much longer barcodes. |
| DeepBarcoding | Yang et al. (2022) | CNN-based classifier. | No code available with publication. |
| mkLTG | Meglécz (2023) | BLAST-based, using thresholds or LCA to avoid over-classification. | Constrained to NCBI taxonomy, output restricted to family, genus, and species. |
| Ensemble NLP | Yadav et al. (2023) | Ensemble NLP & Multinomial Classifier. | No code available with publication. |
| BarcodeBERT | Arias et al. (2023) | Transformer-based architecture pre-trained on unlabeled sequences. | |
| Protax-GPU | Li et al. (2024) | Different implementation of same model as FinPROTAX. | No instructions for training on new data available. |

Table S2: Origin of the backbone trees used to build the phylogenetically constrained reference tree.

| Taxon | Citation | Tree | Additional information |
|---|---|---|---|
| Coleoptera | McKenna et al. (2019) | Suppl. fig. S10 | Dataset_S01: Table S6 |
| Lepidoptera | Kawahara et al. (2019) | Suppl. fig. S12 | |
| Diptera | Wiegmann et al. (2011) | Suppl. fig. S1 | Suppl. info st01 |
| Hymenoptera | Blaimer et al. (2023) | Figure 2 (file in 2.2.2/HYM-mcmctree-70%-topC1.tre) | |
| Trichoptera | Ge et al. (2023) | Align_matrices_trees/85species/5-tree/PCG12_RNA/BI_bpcomp.con | |
| Odonata | Kohli et al. (2021) | Figure 1 | |
| Neuroptera, Megaloptera | Winterton et al. (2018) | Figure 1 | |
| Neuroptera: Myrmeleontidae | Machado et al. (2019) | | |
| Neuroptera: Chrysopidae | Winterton et al. (2019) | | |
| Orthoptera | Song et al. (2020) | Suppl. Arch. 3/dated_trees/FigTree_C80_run1.tre | |
| Ephemeroptera | Tong et al. (2022) | Figure 6 | |
| Plecoptera | Ding et al. (2019) | Figure 6 | |
| Siphonaptera | Zhu et al. (2015) | Suppl. figure S1 | |
| Dermaptera | Wipfler et al. (2020) | Suppl. figure S1 | |
| Mecoptera | Hu et al. (2015) | Figure 1 | |
| Hemiptera, Psocoptera, Thysanoptera | Johnson et al. (2018) | Suppl. Arch. 4/ML_tree_inference_amino_acid/bestTree.newick | |
| Araneae | Garrison et al. (2016) | Figure 2 | |
| Pseudoscorpiones | Benavides et al. (2019) | Figure 4 | |
| Opiliones | Fernández et al. (2017) | Figure 2 | |
| Sarcoptiformes | Klimov et al. (2018) | Suppl. Data 7 | |
| Insecta | Misof et al. (2014) | Figure 1 | |
| Chelicerata | Lozano-Fernandez et al. (2019) | MatrixA_CAT-GTR+G_ReducedTaxon.tre | |
